# Variations in photoreceptor throughput to mouse visual cortex and the unique effects on tuning

**DOI:** 10.1101/2020.11.03.366682

**Authors:** I. Rhim, G. Coello-Reyes, I. Nauhaus

**Author notes:** **Correspondence:** Ian Nauhaus Department of Psychology, Center for Perceptual Systems University of Texas at Austin 108 E. Dean Keeton, Austin TX, 78712.

## Abstract

Visual input to primary visual cortex (V1) depends on highly adaptive filtering in the retina. In turn, isolation of V1 computations to study cortical circuits requires control over retinal adaption and its corresponding spatio-temporal-chromatic output. Here, we first measure the balance of input to V1 from the three main photoreceptor opsins – M-opsin, S-opsin, and rhodopsin – as a function of light adaption and retinotopy. Results show that V1 is rod-mediated in common laboratory settings, yet cone-mediated in natural daylight, as evidenced by exclusive sensitivity to UV wavelengths via cone S-opsin in the upper visual field. Next, we show that cone-mediated V1 responds to 2.5-fold higher temporal frequencies than rod-mediated V1. Furthermore, cone-mediated V1 has smaller RFs, yet similar spatial frequency tuning. V1 responses in rod-deficient (Gnat1^−/−^) mice confirm that the effects are due to differences in photoreceptor contribution. This study provides foundation for using mouse V1 to study cortical circuits.

## INTRODUCTION

The visual system has a cascade of light adaptation mechanisms that accumulate to provide a wide dynamic range. Light adaptation in the retina is first built from differences in rod and cone sensitivity, followed by their respective downstream circuits. In each case, sensitivity increases with a drop in ambient light, which is traded for spatio-temporal-chromatic resolution. That is, monochromatic rods are sensitive to minute changes at the lowest light levels, which are sent through pathways that integrate over a broader window of space and time than cones (Demb and Singer, 2012; Gouras and Link, 1966; Völgyi et al., 2004; Wang et al., 2011). In turn, rods and cones set the stage for subsequent branching of parallel pathways that remain relevant to information processing in the rest of the visual system (Wässle, 2004). To quantify downstream transformations requires knowledge of retinal output, and thus rod vs. cone contributions for a given visual stimulus.

Genetic tools in the mouse have become powerful means to dissect mechanisms of classical functional properties in early visual pathways (Huberman and Niell, 2011), yet cortical studies are based on varying and unknown degrees of input from rods vs. cones. The tendency has been to adopt a similar average light level as that used in classic studies with larger mammals and commercial displays (~ 40 cd/m^2^). On the one hand, there are reasons to believe a similar luminance will place the rodent visual system in a photopic regime, which is based on the anatomy and in-vitro physiology of the retina. The mouse retina has in place the major architectural hallmarks of the primate retina (Wässle and Boycott, 1991). Notably, this includes a substantial majority (>95%) of rods in the photoreceptor mosaic, which then feed into rod bipolar and AII amacrine cells. On the other hand, most primate studies are done near the fovea where the proportion of cones is much higher. Also, the mouse’s eye and pupil are much smaller, which affect retinal irradiance, and thus rod saturation for a given lighting environment. Adding to the challenge of cross-species comparison, most retinal studies on rod saturation performed in-vitro do not include the optics of the eye and the pigment epithelium, making it more difficult to extrapolate to the in-vivo preparation.

Here, we measured V1 responses as a function of graded changes in rod saturation. The intensity and spectral profile of our display is theoretically able to drive rods several fold more than common commercial displays, yet is incapable of saturating rods without artificial pupil dilation. Next, using a simple model that linearly combines the input from three photoreceptor opsins - rhodopsin, cone S-opsin, and cone M-opsin - we quantified the relative contributions of each as a function of photoisomerizations(R*)/rod/sec and retinotopic location. In summary, the model predicts that the dimmest and brightest settings yield 75% and 5% rod input, relative to cones. Our data on percent rod input versus light level also constrained a model of how the cone mosaic’s dorsoventral gradient of M and S-opsin expression are mapped onto the V1 topography. Finally, our calibrated drive of the photoreceptors allowed us to characterize cone- and rod-based spatio-temporal properties in V1 with 2-photon imaging. We showed that spatial tuning changes little between rod- and cone-mediated vision. However, cone-mediated V1 encoded substantially higher temporal frequency than rods.

## RESULTS

### Measurements of photoreceptor contributions to mouse V1

We imaged mouse V1 responses to visual stimuli that were calibrated to modulate distinct photoreceptor opsins across the retina. Two-photon and widefield calcium imaging of neurons was performed after viral-mediated expression of GCaMP6f (Chen et al., 2013a). Figure 1 outlines features of the mouse preparation and visual stimulus paradigm that were common to all results from the study. Most recordings were done in wild-type (WT) mice, and a subset of experiments were repeated in mice lacking rod function (Gnat1^−/−^) (Calvert et al., 2000). The WT mouse retina has 3 main photoreceptor opsins – cone M-opsin, cone S-opsin, and rod rhodopsin – which have unique spectral sensitivities functions (Fig. 1a) and spatial distributions across the retina (Fig. 1b). The upper region of the visual field is filtered primarily through the S-opsin and rhodopsin sensitivity functions. Conversely, in the lower extreme of the visual field, images are filtered through the M-opsin and rhodopsin sensitivity functions (Wang et al., 2011), which are nearly identical.

**Figure 1.**
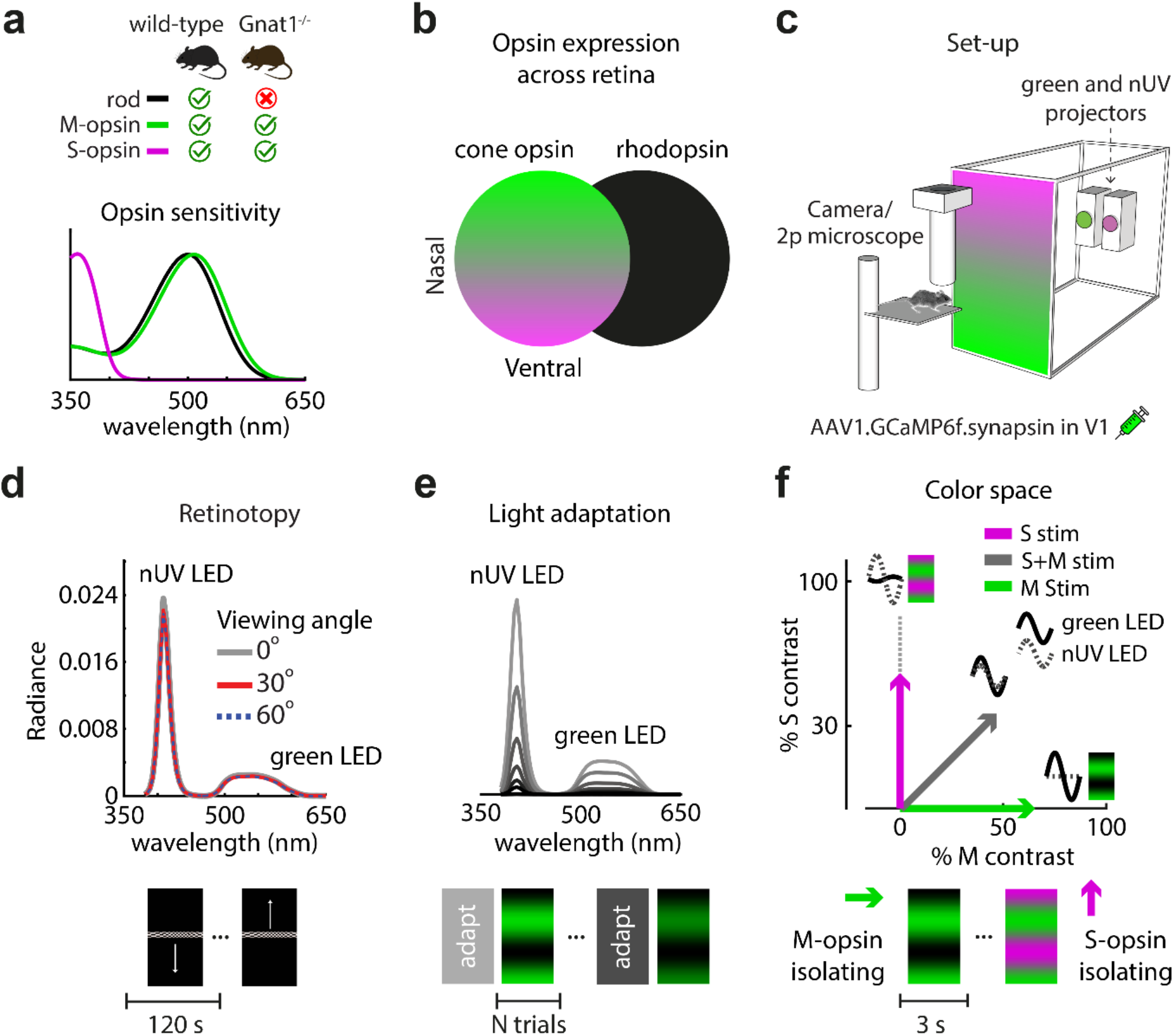
Measurements of photoreceptor contributions to mouse V1. (a) Opsin sensitivity functions for 3 different types of opsins found in rod and cone photoreceptors. Rhodopsin peaks at 498nm, while cone S-opsin and M-opsin peak at 360nm and 508nm, respectively. 2 mouse genotypes were used, wild-type and Gnat1^−/−^. Gnat1^−/−^ is a rod-deficient transgenic strain. (b) Illustration of mouse retina’s cone S/M-opsin expression gradient (left) versus rhodopsin uniform expression (right). Ventral retina (upper visual field) expresses predominantly S-opsin while dorsal retina (lower visual field) expresses predominantly M-opsin. (c) Rear projection of visual stimuli using two monochromatic projectors, near-UV (nUV) and green. Screen color depicts corresponding S/M-opsin expression in mouse’s retina. (d) Measures of the display’s spectral radiance at three viewing angles. Uniformity is required for controlled light adaptation across the retinotopy. At bottom depicts stimuli used to map retinotopy. (e) A total of 6 background light levels were used to adapt the retina, with each level scaling the spectral radiance by a factor of 2. At bottom is a depiction of the adaptation paradigm. Prior to a block of test trials with drifting gratings, the retina was adapted for 10 minutes by the midpoint (i.e. “gray level”) of the test trials. (f) Drifting gratings were calibrated to oscillate along one of three axes in S and M cone-opsin space: S, M, and S+M. The solid and dashed sine-wave insets indicate relative green and nUV LED amplitude and phase used for each of the three color axes. At bottom is a depiction of the stimulus paradigm for measuring color preference, where each trial showed a different color at a given adaptation level.

A major goal of the study was to quantify the balance of input to V1 from all three opsins as a function of visual field location and light adaptation. To this end, visual stimuli systematically varied along 3 dimensions – retinotopy, background light level, and color. The vertical retinotopy was mapped at each pixel (widefield imaging) or neuron (2-photon imaging) based on responses to a monochromatic vertical drifting bar (Kalatsky and Stryker, 2003). The display’s emitted spectral power (radiance) was nearly constant across the range of viewing angles under study (+/− 60°), which allowed for controlled measurements of light adaptation across the retinotopy (Fig. 1d). To alter the state of light adaptation, the display’s spectral power was scaled to a new midpoint for 10 min, followed by a continuous block of full-field drifting gratings that maintained the same midpoint throughout (Fig. 1e). As will be shown, varying the state of light adaptation with our display allowed for a wide dynamic range of rod saturation.

Within each block of light adaptation, drifting gratings were shown at variable orientation to generate robust V1 responses along isolated axes of color space, and/or along a range of spatio-temporal frequencies. Initially, we show results where spatio-temporal frequency is kept constant and responses are compared between M and S cone-opsin contrast (Fig. 1f). These measures of M/S color tuning vs. light adaptation are used to model a) rod vs. cone inputs to V1 as a function of rod photoisomerization rates, along with b) the retinotopic distribution of pure cone-opsin inputs to V1. Later, we show results from varying spatio-temporal frequency, in conjunction with light adaptation and color, in order to compare rod-mediated and cone-mediated tuning properties in V1.

### A bright display does not yield cone-mediated V1 responses if the pupil is not fully dilated

A major goal of this study was to identify a model that allows for predictions of the balance of rod and cone input to V1 across a wide range of background lighting environments. To constrain such a model, our experimental preparation must be capable of saturating the rods to the point of a photopic (i.e. cone dominant) regime. Here, we asked the preliminary question of whether a fully dilated pupil is required to reach a cone-mediated regime with our display, which is much brighter than more commonly used displays. We first swept through 6 levels of light adaptation (b1-b6) without artificial dilation, and then again with dilation using tropicamide (Fig. 2a,b). In each of the 12 blocks of trials, color tuning in the M/S-opsin plane was measured. There is a clear expectation of how color preference changes with increasing rod saturation – as light levels increase, there will be an asymptotic increase in preference for the S-opsin over the M-opsin gratings. In summary, preference for S-opsin approaches much higher values when dilated, showing that cone-mediated vision cannot be achieved without dilation. Below, we describe this main result in finer detail. This same experimental paradigm, but limited to a dilated pupil, is used in later sections to obtain a more detailed model of the balance of photoreceptor inputs across the retinotopy.

**Figure 2.**
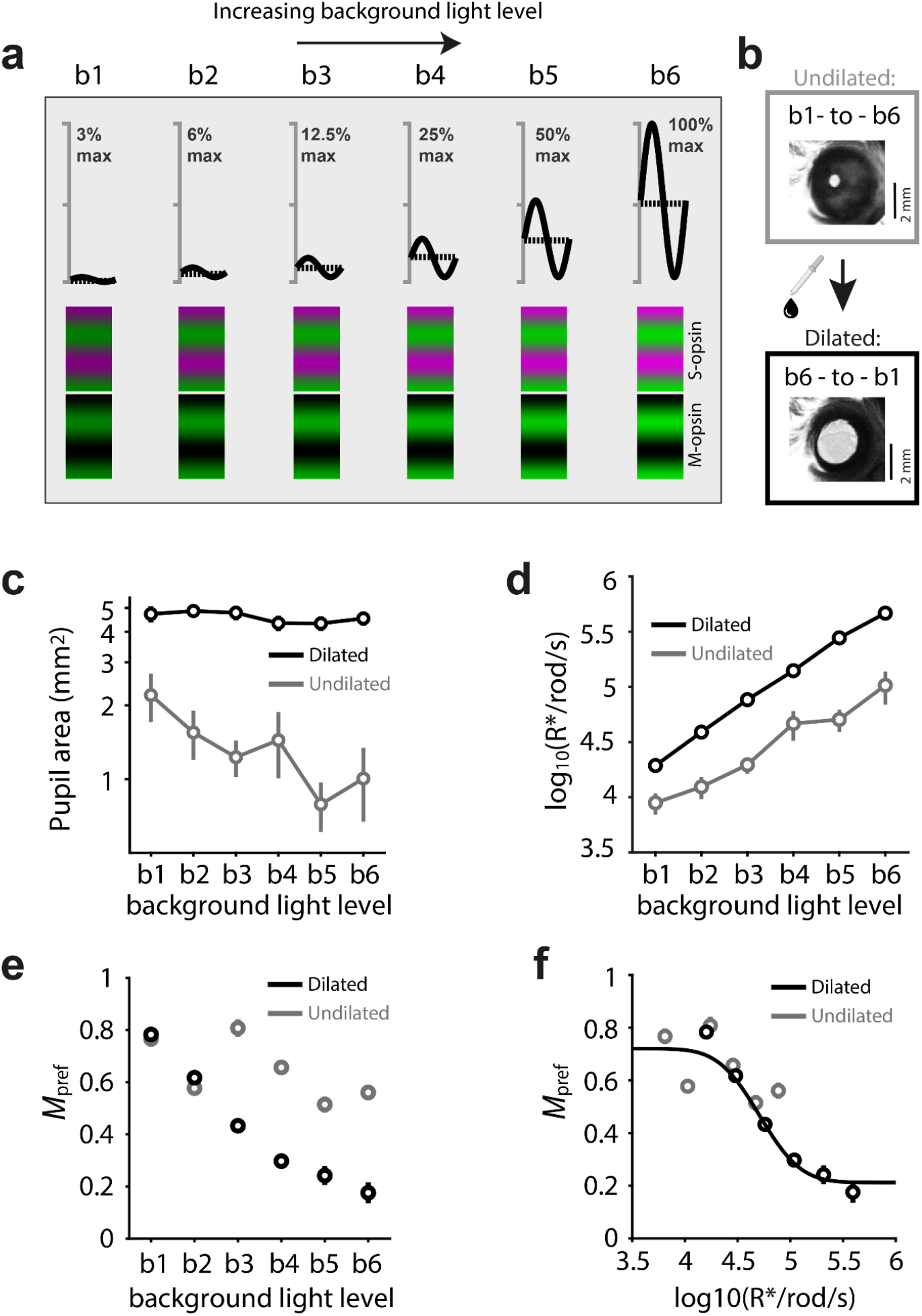
Variable states of retinal adaptation via background light level and pupil diameter. (a) Illustration of the stimuli used to measure color tuning across 6 states of background light adaptation, “b1-b6”. As shown in the top row, adjacent adaptation blocks have mean gray levels that differ by a factor of 2, and constant contrast. Prior to each block, the eye was adapted at the same gray level for 10 min. Bottom two rows: each trial within a block had either a S-opsin (violet and green) or M-opsin (black and green) isolating grating. (b) The experiment began with 6 adaptation blocks, in ascending order (b1-to-b6), shown to an undilated (naive) pupil. Next, the pupil was dilated using tropicamide and reshown the 6 stimulus adaptation blocks in descending order (b6-to-b1). (c) Mean and standard error of pupil area at 6 light levels (x-axis), and the two states of pupil dilation. (d) Same data as in ‘c’, where R*/rod/sec was computed from pupil area and light level. (e) Mean and standard error of preference for the M-opsin over the S-opsin gratings, “*M*_pref_“, for neurons with upper receptive fields (>20° above midline). (f) The same *M*_*pref*_ data points are shown as in ‘e’, but the x-axis was transformed into rod photoisomerization rate, which normalizes for pupil dilation. Sigmoidal curve is fitted to indicate continuity of data between undilated and dilated pupil responses.

Prior to tropicamide administration, pupil size naturally changed across the six background light levels, b1-b6. Specifically, the pupil area was reduced by 2-fold due to the 32-fold scaling in light level. Following administration of tropicamide, the pupil area remained constant, independent of the 6 light levels on the display (Fig. 2c). Using the pupil diameter measurements, light levels were converted into units of rod photoisomerization rates (R*/rod/sec) (Methods), which is a normalized metric for predicting rod saturation from color tuning in each of the adaptation conditions.

For each neuron from the 2-photon imaging, we calculated a metric for color preference, *M*_pref_ = *F*_*M*_/(*F*_*S*_+*F*_*M*_), where *F*_*M*_ and *F*_*S*_ are the mean fluorescence changes in response to the M-opsin and S-opsin stimuli, respectively. *M*_pref_ = 0 indicates that a neuron responds to S gratings, but not to M gratings, with the opposite being true for *M*_pref_ = 1. *M*_pref_ will decay with increasing rod saturation, which is formalized with a model in the Methods (Eq. 5a) and may be intuited as follows. Rhodopsin and M-opsin sensitivity functions are very similar (Figure 1a), so both are preferentially driven by M gratings. Therefore, if rods are the dominant photoreceptor input across the V1 topography, V1 neurons will invariably favor the M-opsin gratings (i.e. greater *M*_pref_). As rods saturate with increasing light level, the expectation is a monotonic decrease in *M*_pref_ that will asymptote to the underlying balance of M and S cone opsin. We limited analyses to the upper visual field, where S-opsin dominates in cones, as this provides a steeper and clearer drop in *M*_pref_ with increasing light levels. This very trend is observed in Figure 2e. A key difference between the dilated and undilated trends in Figure 2e is the relative drop in *M*_pref_ at the highest light levels – they yield comparable *M*_pref_ near b1, but *M*_pref_ falls to much lower values at b6 in the dilated case. We interpret the relatively large *M*_pref_ values at the maximum display intensity (b6) in the undilated case to mean that our display cannot induce rod saturation through an undilated mouse pupil. Our display at b6 has far more rod isomerizing power than commercial displays. For a qualitative reference, most commercial displays have “luminance” values under 100 cd/m^2^, whereas the mean luminance of our green LED at b6 was 360 cd/m^2^. At the same time, luminance ignores the spectral power produced by the nUV LED in our set-up, which will also help saturate mouse rods (Fig. 1a,d). Specifically, the green LED and nUV LED account for 63% and 37% of the rod isomerizing power in the display based on their overlap with the rhodopsin sensitivity function, respectively.

To validate the measurements, we replotted *M*_pref_ as a function of R*/rod/sec, which normalizes for the pupil dilation. Specifically, a line was fit to the two trends in Fig. 2d, which allowed for a conversion of the x-axis in Fig. 2e (b1-b6) into the x-axis of Fig. 2f (R*/rod/sec). As expected, the data in the dilated and undilated conditions now coincide along a similar curve (Fig. 2f). In summary, to push the mouse retina across a wide dynamic range of light adaptation, which is needed to constrain a model, our setup requires artificial dilation to maximize photon flux at the retina.

### Photoreceptor contributions to V1 as a function of isomerization rates and visual field location

Here we describe results from the same visual stimulus paradigm described above, which varies color and light adaptation, but we limited the analyses to the dilated pupil. Since rods are monochromatic and uniformly distributed across the mouse retina, the entire visual field at night is filtered through a common spectral sensitivity function. However, when the dichromatic cone population is allowed to take over, there is a gradient of spectral sensitivity along the retina’s dorsoventral axis that is conveyed to V1 and higher visual areas (Rhim et al., 2017). What are the dynamics of the monochromatic-to-dichromatic photoreceptor transformation at the level of V1 topography, and does our current setup allow for it to be resolved and thus modeled? To address these questions, we used the same experimental paradigm described in Figure 2a but limited to a fully dilated eye; i.e. S- and M-isolating gratings of equal contrast were presented after adapting to 6 different light levels.

Figure 3 shows that the graded unveiling of the color preference map is clearly resolved. At the lowest light levels, neurons across the entire V1 retinotopy were mostly responsive to M-isolating gratings, indicating a predominance of rod input. Increasing light levels induced a graded *exchange* in responsiveness: responses to M-isolating gratings fell, and responses to S-isolating gratings rose. In the upper visual field, the exchange is especially dramatic, as this part of the cone mosaic is dominated by S-opsin – neurons go from strong preference for the M-opsin gratings (*M*_pref_ ~ 1) to strong preference for S-opsin gratings (*M*_pref_ ~ 0) with increasing light level. These examples confirm that the display generates a sufficient range of both intensity and color to model photoreceptor contributions across the V1 topography. Next, the two-photon data from this experiment was combined across animals to model a) the balance of rod and cone inputs as a function of light level and b) the balance of S vs. M cone-opsin inputs as a function of retinotopy.

**Figure 3.**
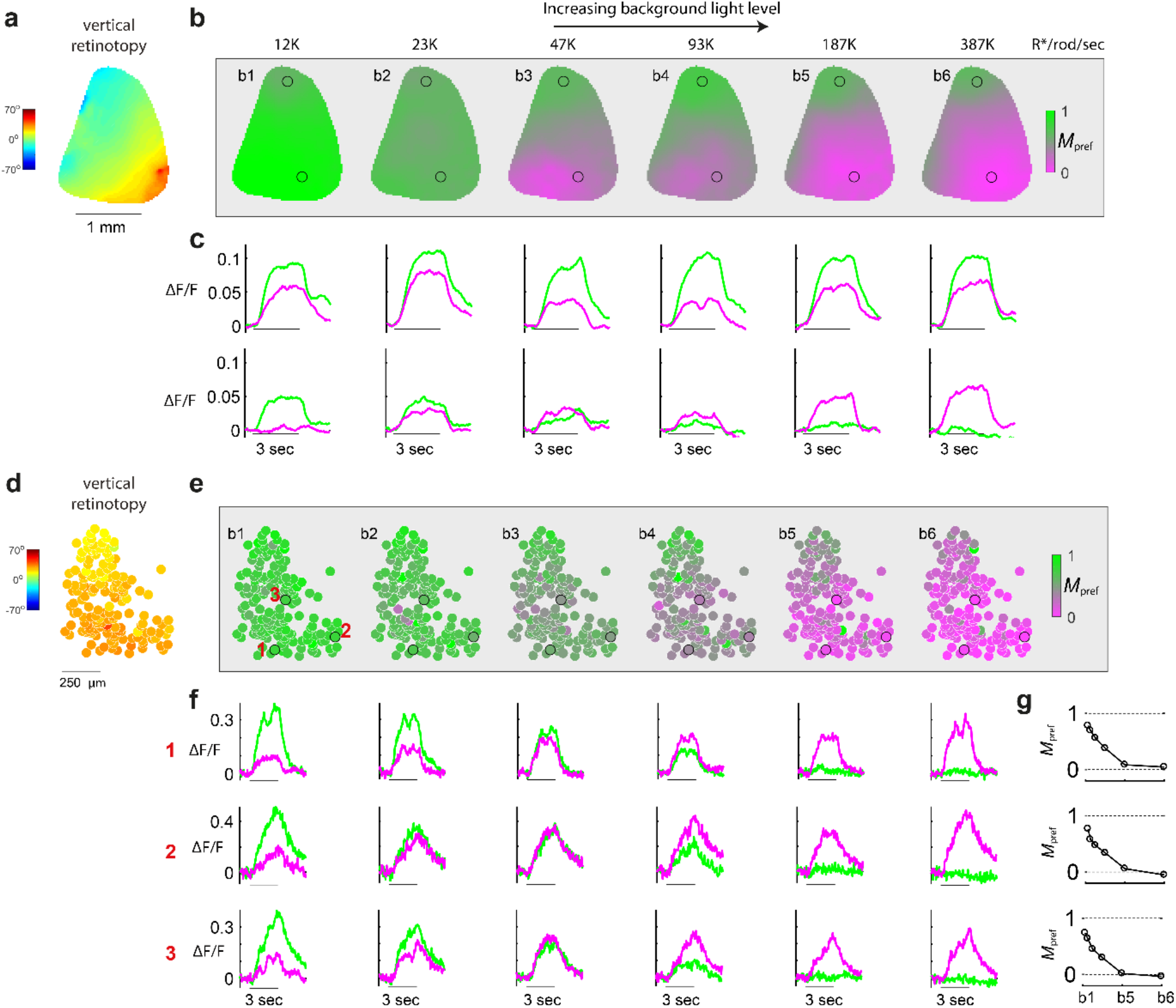
Revealing the cone mosaic’s M-to-S gradient in V1 via progressive rod saturation. (a-c) Wide-field calcium imaging. (a) Map of V1’s vertical retinotopy. (b) Maps of *M*_pref_, at 6 states of light adaptation. *M*_pref_ at each pixel indicates the preference for the M-opsin color axis, over the S-opsin color axis. At low light levels, *M*_pref_ ~1 at all pixels, indicating rod input. At higher light levels, rods saturate, revealing the M-to-S cone opsin gradient. (c) Time courses of fluorescence signal at the 2 ROI’s indicated in the map. Top circle corresponds to lower visual field; bottom circle corresponds to upper visual field. Green and violet traces are responses to M-opsin and S-opsin gratings, respectively. The line under each pair of traces indicates the stimulus duration. (d-g) Two-photon imaging. (d) V1 map of vertical retinotopy, where each circle is the location of a neuron. (e) Maps of *M*_pref_, at 6 states of light adaptation. (f) Each row shows time courses of a different neuron (see their locations in *M*_pref_ map at b1). Green and violet traces are responses to M-opsin and S-opsin gratings, respectively. Right-most column shows each neuron’s mean *M*_pref_ during stimulus presentation, as a function of light level.

#### Model description

Next, we describe key elements of the model used to quantify photoreceptor contributions to V1. The equations given in the Methods describe the response of a V1 neuron to each chromatic grating (S and M-opsin direction) as a linear combination of 3 inputs: rods, S-opsin, and M-opsin. The magnitude of rod inputs depends only on light levels (not retinotopy), whereas the magnitude of S and M cone opsin inputs depends only on location along the vertical retinotopy (not light level). The first assumption here is that rod inputs are uniform across the visual field in an animal that lacks a fovea. The second assumption is that cone responses maintain Weber adaptation, meaning that responses depend on contrast and not the range of background light levels used in this study. We tested this latter assumption, given the proximity of our lowest light level to cone thresholds (i.e. their “dark light”) established in prior studies (Naarendorp et al., 2010; Schnapf et al., 1990).

To do this, we measured V1 responses as a function of light level in mice with dysfunctional rods (Gnat1^−/−^) (Calvert et al., 2000), which showed that they do not increase with light level, for either S or M-opsin gratings (Fig 4a). Figure 4a uses cells at all locations of the retinotopy, whereas Figure 4b limits the population to neurons with RFs in the upper region of the visual field (> 30° above midline). In both cases, responses do not change significantly with light level (p > 0.05; Pearson correlation). Figure 4b also shows that neurons in the upper visual field have near zero response to the M-opsin gratings, as expected. The assumption of Weber adaptation for cones simplified the model, yet accounts for a wide range of lighting conditions. The model can be summarized below using the unitless metric *M*_pref_, which again is the response to the M-opsin gratings, divided by the sum of responses to the S-opsin and M-opsin gratings.

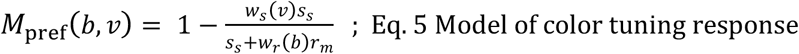

**Figure 4.**
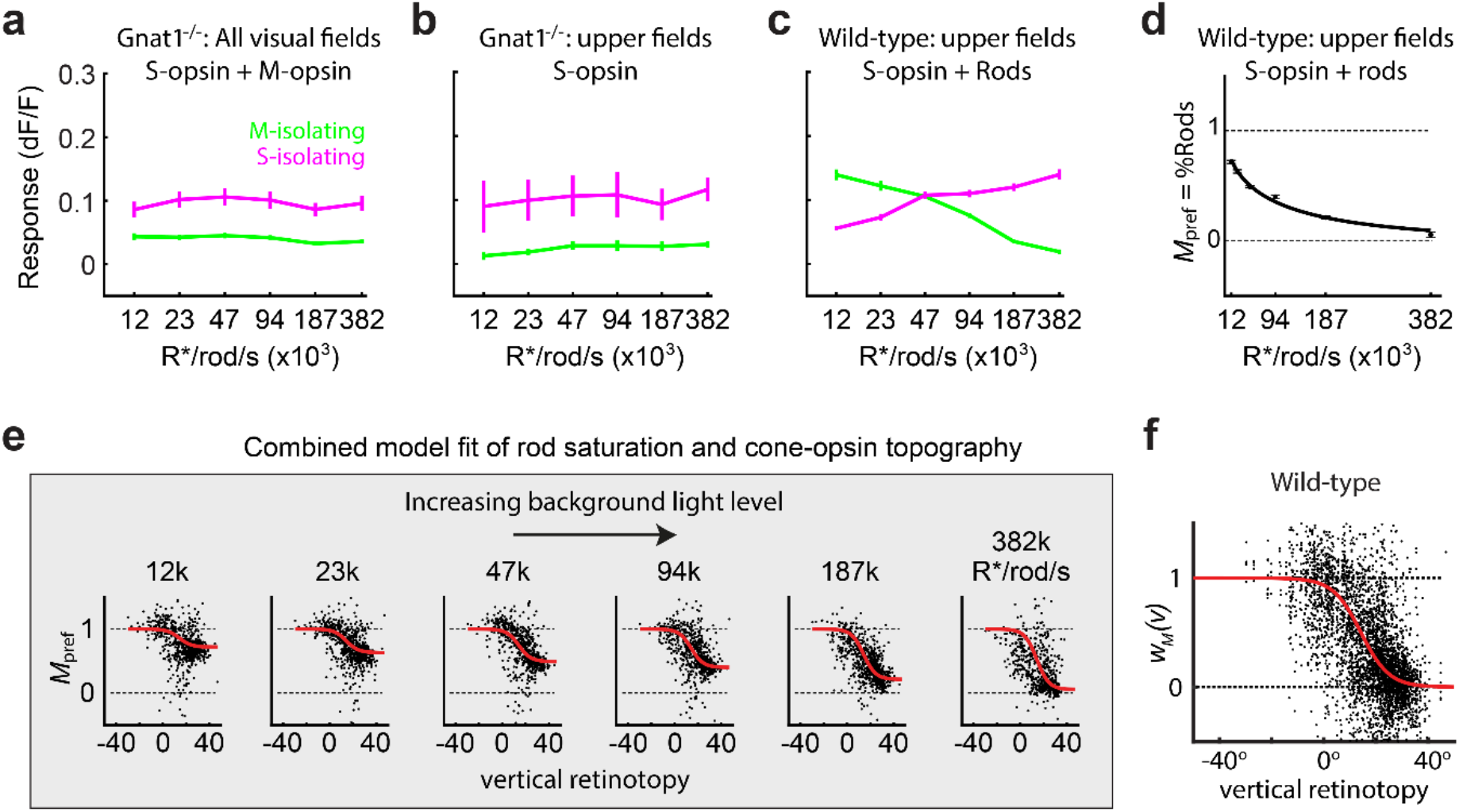
Modeling photoreceptor inputs as a function of light adaptation and vertical retinotopy. (a) Mean and standard error responses of V1 neurons to the two different color axes, in Gnat1^−/−^ mice, at 6 light levels. This was used to confirm that cones are in a constant state of Weber adaptation across the range of light levels. (b) The same analysis was performed as in ‘a’ using only upper visual field neurons, which limits the cone inputs to S-opsin. (c) Mean and standard error responses of V1 neurons in WT mice, at 6 light levels. Data was limited to the upper visual field to highlight the population’s switch in responsiveness from rod-to cone-mediated inputs. (d) The color-tuning metric, *M*_pref_, when limited to neurons in the upper visual field of WT mice, is effectively a measure of the balance of rod vs. cone input, %Rods. The mean and standard error bars of %Rods are shown as a function of light level, and the line shows the model fit. (e) The six panels of *M*_pref_ vs. vertical retinotopy show data points from the same WT population, at the six states of light adaptation. Overlayed in red is the model fit of *M*_pref_ as a function of vertical retinotopy and light adaptation (see Methods). (f) The y-axis, *w*_*m*_*(v)*, is an estimate of the balance of M- and S-cone opsin providing input to V1 at each retinotopic location. 1 and 0 indicate 100% M- and S-opsin, respectively. Data points are an accumulation from all the panels in ‘e’, after normalizing by the model fit along the dimension of light intensity using the %Rod model in ‘d’. The red line is the model fit of the retina’s cone opsin gradient (M vs. S cone-opsin input) vs. vertical retinotopy, recapitulated in V1.

The two functions we are trying to solve for are *w*_*r*_(*b*) and *w*_*s*_(*v*). *w*_*r*_(*b*) is the unitless weight of rod input as a function background light level. *w*_*s*_(*v*) is the unitless weight of S-opsin input as a function of vertical retinotopy. The corresponding weight of M-opsin input is simply *w*_*M*_(*v*) = 1 − *w*_*s*_(*v*). *S*_J_ is the S-opsin contrast of S-opsin gratings (~0.6), and *r*_*m*_ is the rod contrast of the M-opsin gratings (~0.6). Under complete rod saturation, *w*_*r*_(*b*) approaches zero, giving *M*_pref_(*b*, *v*)~ 1 − *w*_*s*_(*v*) = *w*_*m*_(*v*), which is the pure cone opsin gradient. Oppositely, under low light levels, *w*_*s*_(*v*) ≪ *w*_*r*_(*b*), giving *M*_pref_(*b*, *v*)~ 1, which is pure rod input that only responds to the M-opsin gratings across the entire V1 topography. Below, we describe the results of fitting *w*_*s*_(*v*) and *w*_*r*_(*b*), based on the measurements of *M*_pref_(*b*, *v*).

#### Model fit of rod saturation

To fit the model to data, we used two-photon responses from 11 WT mice. Figure 4e plots *M*_pref_ against vertical retinotopy, at all 6 light levels. The population shows graded rod saturation and the emerging cone mosaic with increasing light levels – i.e. from left-to-right, *M*_pref_ is reduced and becomes more steeply correlated with the retinotopy. To simplify the relationship between rod saturation and *M*_pref_, we limited the data to the subpopulation of neurons that represent the upper visual field (> 30°), which limits the photoreceptor opsin input to cone S-opsin and rods. Studies in the mouse retina show that cone signals in the upper visual field are almost exclusively mediated by S-opsin (Applebury et al., 2000; Wang et al., 2011) – i.e. *w*_*s*_(*v* > 30) = 1, where 30° is a conservative threshold based on these prior studies. By limiting responses to the upper field, it can be shown that the measured color preference, *M*_pref_, is equal to the proportion of rod input, relative to cones, defined as %Rod(*b*) = *w*_*r*_(*b*)/[*w*_*r*_(*b*) + *w*_*s*_(*v* > 30)]. In turn, the y-axis of Fig. 4d is shown as “*M*_pref_ = %Rod”. The trend of %Rod(*b*) is well-fit by the equation 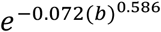. The data shows that the minimum and maximum light level of the display (b1 and b6) yield rod contributions to V1 of 75% (%Rod = 0.75) and 5% (%Rod = 0.05), relative to the cones.

#### Model fit of M-to-S cone opsin topography in V1

In a previous study, we showed that the M-to-S cone opsin gradient is recapitulated in V1 and higher visual areas (Rhim et al., 2017). However, this prior study did not account for potential rod “contamination” in measuring the cone opsin map, and it employed intrinsic signal imaging. Here, we were able to remove the rod contamination using the model of %Rod(*b*) in Fig. 4d. The topographic map of cone opsin inputs to V1, *w*_*m*_(*v*) = 1 − *w*_*s*_(*v*), is dependent on known variables in our model: 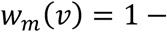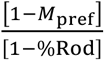. Specifically, this equation converts the measured values of color preference, *M*_pref_(*b*, *v*) (Fig. 4e, all panels), into the cone-opsin topographic map, *w*_*m*_(*v*) (Fig. 4f, black), by way of the model fit to %Rod(b) (Fig. 4d). The fitted red curve in Figure 4f is 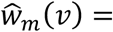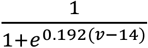. Finally, 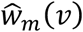 was converted to 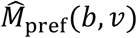, by way of %Rod(*b*), and overlaid on the data points in Figure. 4e (red).

### V1 responds to higher temporal frequencies when stimuli drive cones more than rods

Rods and cones are routed through different pathways in the retina that filter unique spatio-temporal properties of the visual scene for further processing in the cortex. To date, studies of detailed spatio-temporal tuning in mouse V1 are largely rod-mediated, as strongly suggested by the results in Fig. 2 above – in the final section of the Results, this conclusion is further quantified with a detailed simulation. Here, we asked if V1’s spatio-temporal tuning properties change when we vary %Rods - the balance of rod and cone input. Two methods were used to alter %Rods – one that varied the state of light adaptation (i.e. rod saturation), and the other that varied color (i.e. rod and cone contrast). For the light adaptation method, rod-deficient (Gnat1^−/−^) mice were used as a control.

Temporal frequency tuning was measured under two background light levels, b1 (12K R*/rod/sec) and b5 (200K R*/rod/sec), where the predicted %Rod input is 75% (i.e. 25% cones) and 20% (i.e. 80% cones), respectively. Figure 5a-c and Table 1 show that there is a shift in tuning toward higher temporal frequencies when cones provide the primary input. For each neuron, we used two parameters from Gaussian fits to quantify tuning at each level of rod saturation, center-of-mass (CoM_b1_ & CoM_b5_) and the high-pass cut-off frequency (HPCO_b1_ & HPCO_b5_). To quantify differences, we calculated the geometric mean of the ratio between tuning parameters. In turn, paired t-tests were performed on the logs – e.g. log(CoM_b1_) vs. log(CoM_b5_). The geometric mean of CoM_b5_/CoM_b1_ was 1.12 (i.e. 12% change) and was significantly greater than one (p = 2.47e^−35^). The geometric mean of HPCO_b5_/HPCO_b1_ was 1.38 (p = 1.76e^−32^). To verify that these results were due to an exchange in activity between rods and cones, we repeated these experiments and analyses in Gnat1^−/−^ mice (Fig. 5d-f). While the difference in tuning for spatial or temporal frequencies between the two light levels were statistically significant, there were nominal difference in calculated means: geometric mean of CoM_b5_/CoM_b1_ = 1.00 (p = 0.012); geometric of HPCO_b5_/HPCO_b1_ = 1.01 (p = 0.0047).

**Figure 5.**
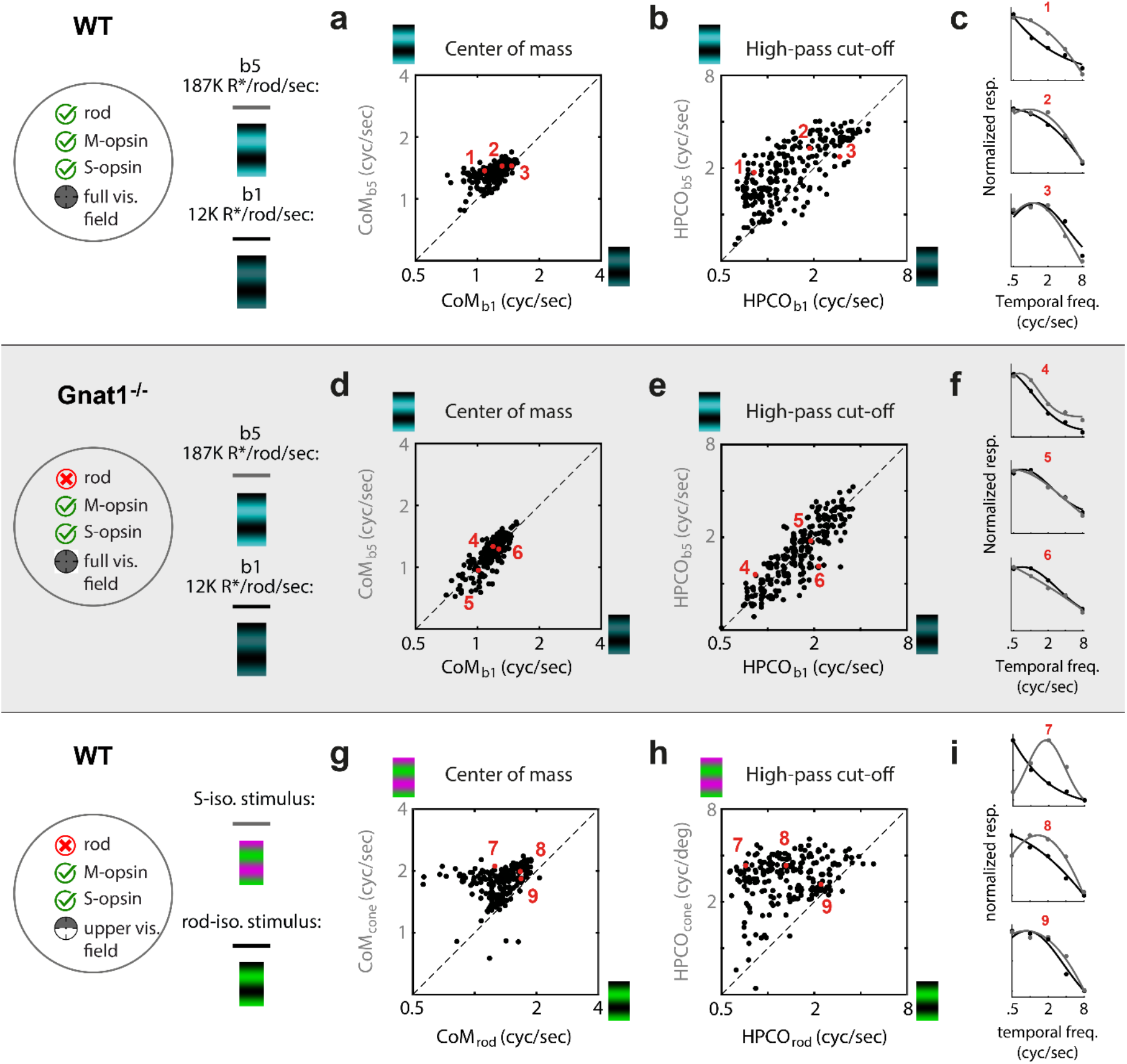
V1 responds to higher temporal frequencies when stimuli drive cones more than rods. Top row (‘a-c’) is data from WT mice, comparing temporal frequency tuning at two levels of rod saturation, b1 and b5 (n=240). Scatter plot compares the temporal frequency tuning curve’s center-of-mass (CoM) at b5 (y-axis) and b1 (x-axis). Unity is dashed black line. The geometric mean of CoM_b5_/CoM_b1_ is 1.12 (i.e. 12% change; t-test: p=2.47e^−35^). (a) Same as in ‘a’, but the temporal frequency tuning parameter is the high-pass cutoff frequency (HPCO). The geometric mean of HPCO_b5_/HPCO_b1_ is 1.38 (t-test: p=1.76e^−32^). (c) Tuning and fits of 3 example neurons at the 5^th^, 50^th^, and 95^th^ percentile of the HPCO_b5_/HPCO_b1_ distribution from ‘b’. Dots are normalized responses and lines are the fits. In each case, black and gray correspond to ‘b1’ and ‘b5’, respectively. Next, middle row of panels (‘d-f’) shows data from rod-deficient Gnat1^−/−^ mice, using the same experiment and analysis as in the top row (n=249). (d) The geometric mean of CoM_b5_/CoM_b1_ is 1.00 (t-test: p=0.012). (e) The geometric mean of HPCO_b5_/HPCO_b1_ is 1.01 (t-test: p=0.0047). (f) Tuning and fits of 3 example neurons at the 5^th^, 50^th^, and 95^th^ percentile of the distribution of HPCO_b5_/HPCO_b1_ from ‘e’. Bottom row of panels (‘g-i’) shows data from WT mice, limited to neurons with a receptive field 30-deg above the retina’s midline where opsin expression is mostly limited to S-opsin and rhodopsin (n=216). Here, background light levels were held constant at b4 or b5 and tuning was compared between stimuli of isolating cone- and rod-contrast. (g) Scatter plot compares temporal frequency tuning CoM between the rod (x-axis) and cone (y-axis) isolating contrasts. Geometric mean of CoM_cone_/CoM_rod_ is 1.42 (t-test: p=1.91e^−31^). (h) Same as in ‘g’, but the parameter measurement for each neuron and light level is HPCO. Geometric mean of HPCO_cone_/HPCO_rod_ is 2.54 (t-test: p=1.97e^−51^). (i) Tuning and fits of 3 example neurons at the 5^th^, 50^th^, and 95^th^ percentile of the distribution of HPCO_cone_/HPCO_rod_.

**Table 1.**
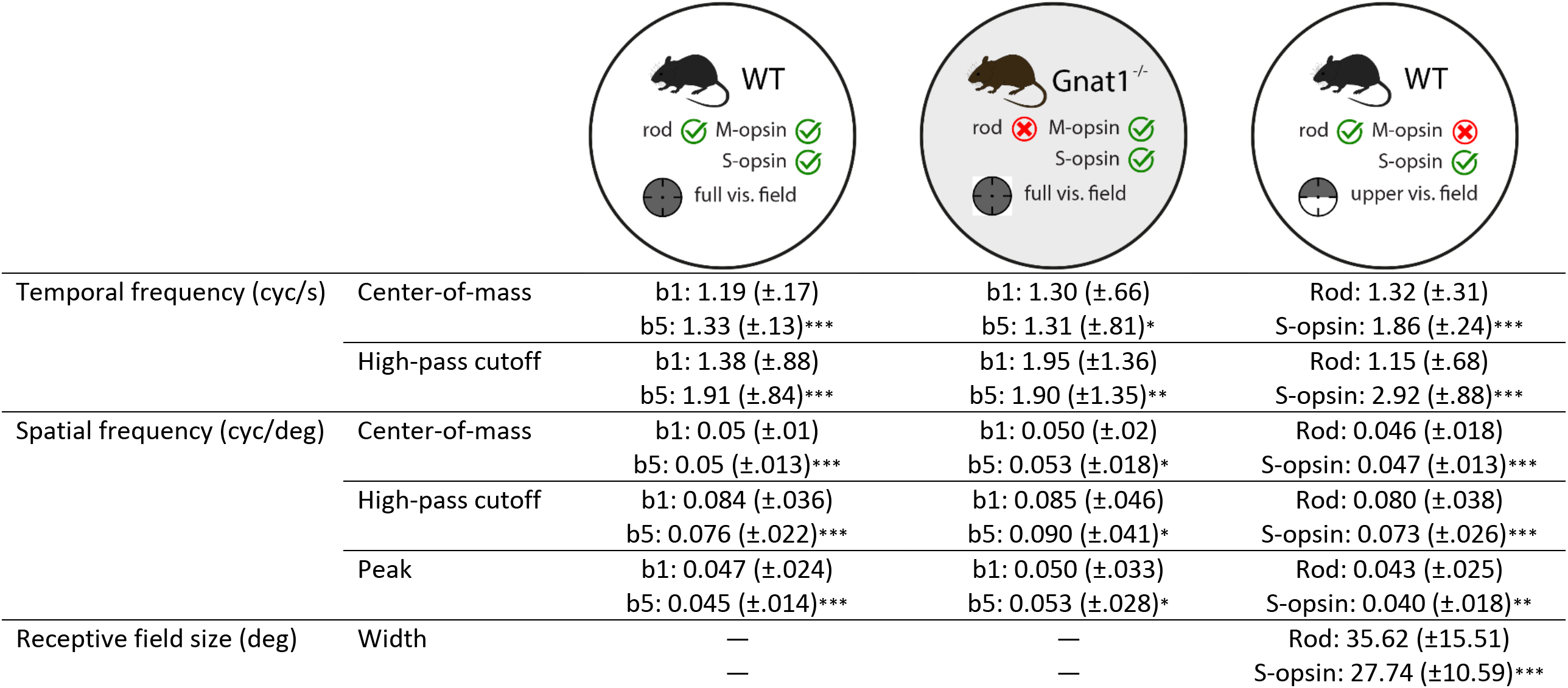
Summary of spatio-temporal tuning estimates, outlining parameter mean and standard deviation values from figures 5–7. Parameter estimates for temporal frequency, spatial frequency, and receptive field size mapping experiments from Figures 5–7, respectively, compiled in table format. First 2 rows outline temporal frequency curve fit parameter estimates of center-of-mass and high-pass cutoff for “b1” (12K R*/rod/sec) vs. “b5” (187K R*/rod/sec) conditions in wild-type mice (3^rd^ column), “b1” vs. “b5” in rodless (Gnat1^−/−^) mice (4^th^ column), and “b5” cone vs. rod inputs in wild-type mice (5^th^ column). “Rod” and “S-opsin” in last (5^th^) column indicate M-opsin contrast and S-opsin contrast stimuli used, respectively. Next 3 rows outline spatial frequency curve fit parameter estimates of center-of-mass, high-pass cutoff, and peak for “b1” vs. “b5” conditions in wild-type mice (3^rd^ column), “b1” vs. “b5” but in rodless (Gnat1^−/−^) mice (4^th^ column), and “b5” cone vs. rod inputs in wild-type mice (5^th^ column). Last row outlines receptive field size comparison, measured as 2σ of Gaussian fit, between “b5” rod vs. cone inputs in wild-type mice. For each measurement listed, “*mu (±std)*” format is used to denote geometric mean value followed by standard deviation for temporal-spatial frequency parameters. “*mu (±std)*” format is used to denote arithmetic mean value followed by standard deviation for receptive field size estimates. “*/**/***” state statistical significance of p-values under .05, .01, and .001, respectively.

Next, we used a different method to compare temporal frequency tuning between rod and cone-mediated inputs. This method kept the light adaptation at a constant mesopic level, but varied rod and cone contrast. Just as in previously described experiments (Figs. 2–4), stimuli had color contrast that isolate either S-opsin or M-opsin (Fig. 1g). However, here the analysis was limited to neurons with upper receptive fields, so the M-opsin stimulus is effectively a rod-isolating stimulus; i.e. M-opsin and rods have nearly identical spectral sensitivity, yet the upper fields lack M-opsin. For the same reason, the S-opsin stimulus is effectively a cone-isolating stimulus (more specifically, the calculated rod contrast is 2% in response to the S-opsin stimulus). Figure 5g-i shows the comparison of temporal frequency tuning between the two different color directions. The results are consistent with the changes induced by different light adaptation levels in Figure 5a-c, in that the cone-dominated tuning is shifted to higher values of temporal frequency. However, the differences are stronger with the method of rod- and cone-isolating contrasts, which seems due to much purer separation of rod and cone drive. The geometric mean of CoM_cone_/CoM_rod_ is 1.42 (i.e. 42% change), and significantly greater than 1 (p = 1.91e^−31^). The geometric mean of HPCO_cone_/HPCO_rod_ exhibited the strongest differential, at 2.54 (p = 1.97e^−51^).

### Cone- and rod-mediated V1 responds to a similar band of spatial frequencies

Similar analyses were performed for spatial frequency tuning as those described above for temporal frequency tuning. Three parameters were taken from Gaussian fits to the spatial frequency tuning curves: peak location, CoM, and HPCO. The comparisons between two levels rod saturation (b1 and b5) in WT mice are shown in Fig. 6a-d and Table 1. The tuning is quite similar between the two states of adaptation. However, there is a small but significant shift in tuning to lower spatial frequencies when cones are the primary input: Geometric mean of peak_b5_/peak_b1_ = 0.94 (p = 2.07e^−7^), CoM_b5_/CoM_b1_ = 0.96 (p = 1.04^−6^), and HPCO_b5_/HPCO_b1_ = - 0.90 (p = 2.4e^−17^). For Gnat1^−/−^ mice, this analysis also revealed a significant but minute difference in tuning for all of the three parameters. However, in this case, the higher light levels yielded tuning for higher spatial frequencies. (Fig. 6e-h, Table 1)

**Figure 6.**
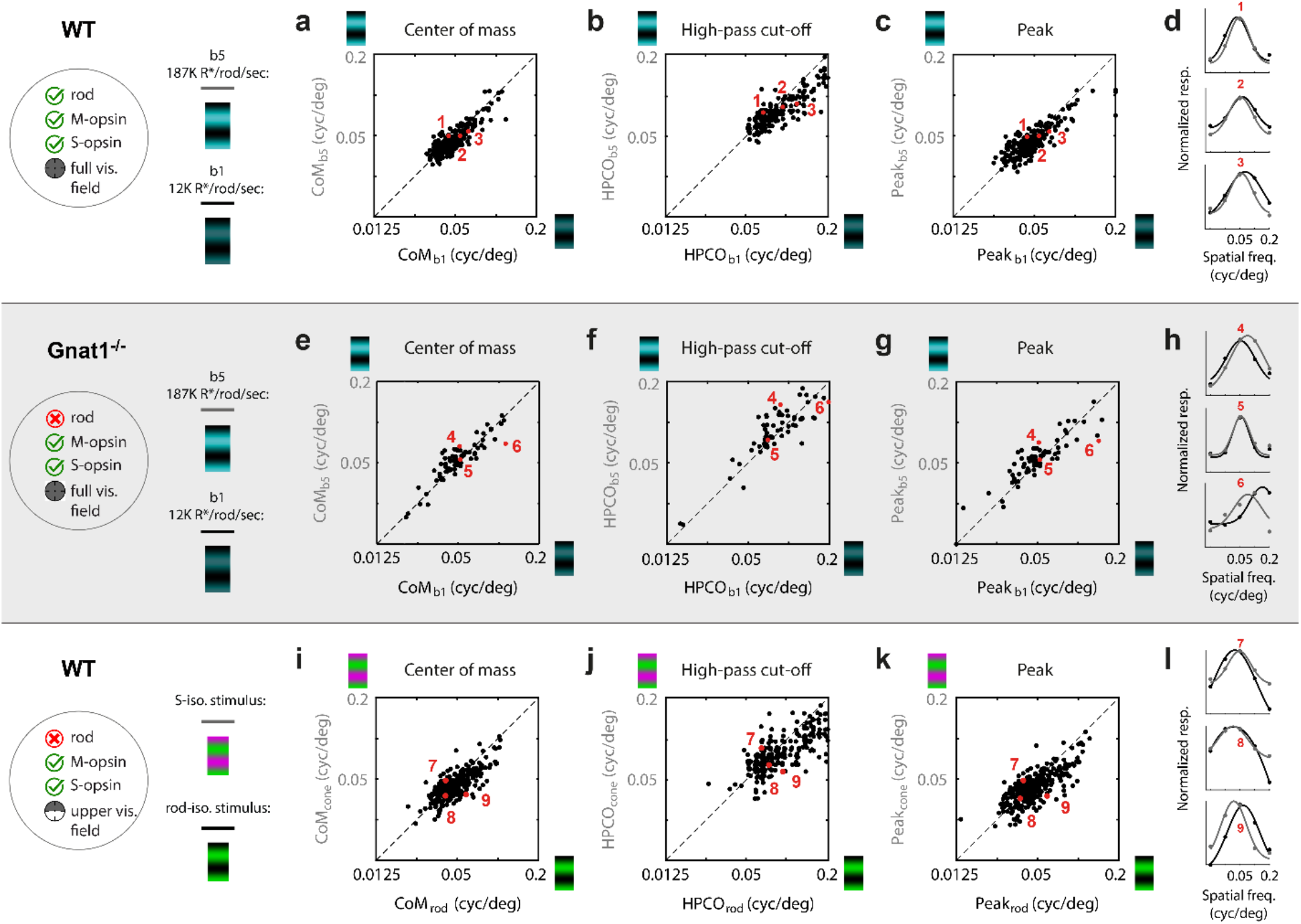
Rod-driven responses in V1 have marginally higher spatial frequency tuning than cone-driven responses. Top row (‘a-d’) shows data from WT mice, comparing spatial frequency tuning at two light adaptation levels, b1 and b5, corresponding to 12K and 187K R*/rod/sec (n=247). (a) Scatter plot compares center-of-mass (CoM) estimates of spatial frequency tuning curve at b5 (y-axis) and b1 (x-axis). Each data point is from a single neuron. Unity line is dashed black. The geometric mean of CoM_b5_/CoM_b1_ is 0.96 (t-test: p=1.04e^−6^). (b) Same as in ‘a’ but the tuning parameter is the high-pass cutoff frequency (HPCO). The geometric mean of HPCO_b5_/HPCO_b1_ is 0.90 (t-test: p=2.40e^−17^). (c) Same as in ‘a,b’ but the tuning parameter is the peak spatial frequency (Peak). The geometric mean of Peak_b5_/Peak_b1_ is 0.94 (t-test: p=2.07e^−7^). (d) Tuning and fits of 3 example neurons at the 5^th^, 50^th^, and 95^th^ percentile of the distribution of HPCO_b5_/HPCO_b1_. Dots are normalized responses and lines are the fits. Black and gray correspond to ‘b1’ and ‘b5’, respectively. Next, middle row of panels (‘e-h’) is data from rod-deficient Gnat1^−/−^ mice, using the same experiment and analysis as in the top row (n=71). (e) The geometric mean of CoM_b5_/CoM_b1_ is 1.05 (t-test: p=.013). (f) The geometric mean HPCO_b5_/HPCO_b1_ is 1.06 (t-test: p=0.021). (g) The geometric mean of Peak_b5_/Peak_b1_ is 1.06 (t-test: p=0.029). (h) Tuning and fits of 3 example neurons at the 5^th^, 50^th^, and 95^th^ percentile of the distribution of HPCO_b5_/HPCO_b1_ from ‘f’. Bottom row of panels (‘i-l’) shows data from WT mice, limited to neurons with a receptive field 30-deg above the midline where expression is mostly limited to S-opsin and rhodopsin (n=366). Here, the background light levels were held constant at b4 or b5 and tuning was compared between stimuli of isolating cone- and rod-contrast. (i) Scatter plot compares spatial frequency tuning CoM between the rod (x-axis) and cone (y-axis) isolating contrasts. Geometric mean of CoM_cone_/CoM_rod_ is 1.02 (t-test: p=3.35e^−4^). (j) Same as in ‘i’ but for HPCO. The geometric mean of HPCO_cone_/HPCO_rod_ is 0.91 (t-test: p=1.79e^−4^). (k) Same as in ‘i’ but for Peak. The geometric mean of Peak_cone_/Peak_rod_ is 0.93 (t-test: p=4.69e^−3^). (l) Spatial frequency tuning and fits of 3 example neurons at the 5^th^, 50^th^, and 95^th^ percentile of the distribution of HPCO_cone_/HPCO_rod_.

Finally, at a mesopic light level, we compared spatial frequency tuning between cone and rod stimuli in the upper visual field (Fig. 6i-l). Similar to the results above with rod saturation, there was not much visually discernable difference in tuning across the two sets of gratings, yet rod mediated V1 responds to significantly higher spatial frequencies. The main findings of spatio-temporal tuning comparison between rod and cone inputs are summarized in Table 1.

### V1 has narrower receptive fields when stimuli drive cones more than rods

To further characterize rod vs. cone-mediated difference in V1’s spatial tuning, receptive field width was measured along the vertical dimension of the retinotopy. Here, to generate stimuli with rod and cone drive, we did not use the method of variable rod saturation (i.e. light level) like in Figures 6a-c and 7a-c. Instead, results are limited to the method of exploiting the unique spectral sensitivity of rods and cones encoding the upper visual field. The visual stimulus consisted of flashed horizontal bars, randomized for color-contrast and vertical position (Fig. 7a). The color-contrast of the bar against the background was along either the S- or M-axis of cone opsin color space.

**Figure 7.**
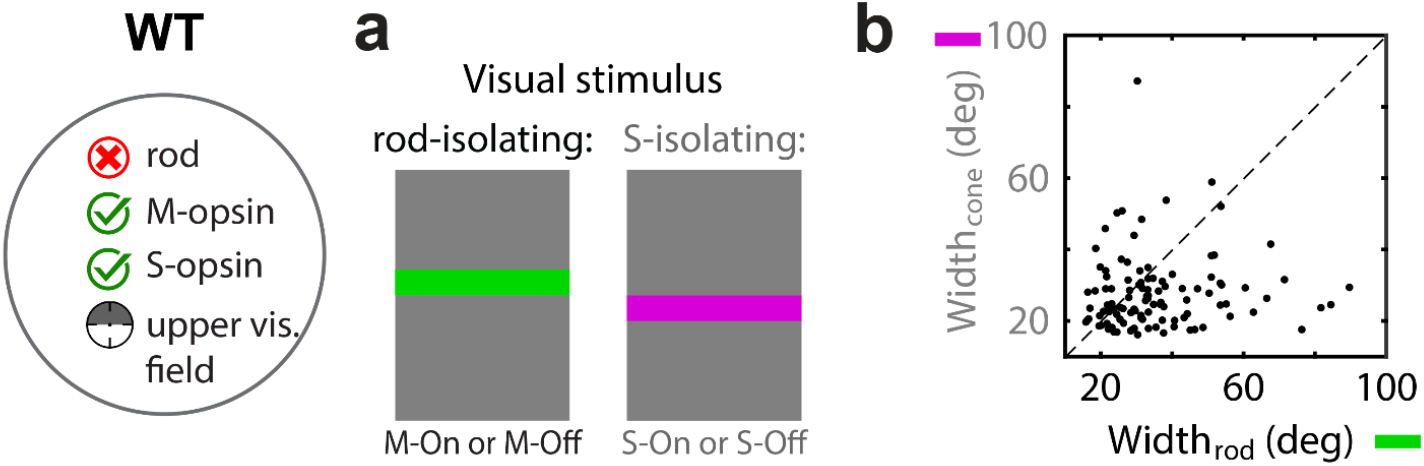
Rod-dominant input to V1 has larger receptive field sizes than cone-dominant input. (a) Horizontal bars (6° in vertical width) sampled the screen at 2.8° intervals. They were flashed intermittently, varying in color (M-ON, M-OFF, S-ON, or S-OFF) at constant background light level at ‘b5’. Data is shown for WT mice, limited to neurons with a receptive field 30-deg above the midline where expression is mostly limited to S-opsin and rhodopsin (n=108). A Gaussian curve is fit to each neuron’s tuning curve of response vs. bar position, for rod and S-opsin contrast, which was used to compare receptive field size. Rod isolating contrasts combine responses from both M-ON and M-OFF bars. Cone isolating contrasts combine responses from both S-ON and S-OFF bars. (b) Scatter plot compares receptive field width (2σ between rod (x-axis) and cone (y-axis) isolating contrasts. Each data point represents a single neuron. Unity line is dashed black. The mean difference is 7.9 visual angle degrees; geometric mean of Width_cone_/Width_rod_ = 0.83 (t-test: 5.04e^−4^).

As described above (Figs. 6g-i and 7i-l), under mesopic adaptation, the S- and M-opsin stimuli in the upper visual field isolate cones and rods, respectively. This yielded two spatial tuning curves for each neuron, to which the σ of Gaussian fits quantified the receptive field width under cone- and rod-mediated inputs (Figure 7; Table 1). The mean receptive field width (2σ) for rod- and cone-mediated inputs was, 35.6° and 27.7°, respectively. The average difference was 7.9°, with geometric mean of RF_S-opsin_/RF_Rod_ = 0.83 (p=5.04e^−4^; t-test). Wider V1 receptive fields when rod-mediated is consistent with an inheritance from subcortical receptive fields (Cowan et al., 2017; Grimes et al., 2014). It may also seem at odds with the results in Figure 6 on spatial frequency tuning, but as described in the Discussion, it is consistent with a simple model of how V1 receptive fields arise from geniculate inputs.

### %Rod inputs to mouse V1 when exposed to outdoor lighting and commercial displays

In Figure 4d, we used responses to variable light intensity to fit a model of the percentage of rod input to V1, relative to cone input, as a function of rod isomerization rate; viz. %Rod(*b*). Here, we used this model to make predictions of %Rod under two other relevant environments. We first asked what %Rod values are achieved under “standard” luminance (cd/m^2^) intensity produced by a commercial display. We show that these standard luminance levels yield mostly rod input. We then asked if the mouse retina can achieve a photopic state when exposed to natural outdoor lighting, and if so, what time of day the transition occurs.

Commercial displays do not drive cone S-opsin, but do drive rods and cone M-opsin. To make predictions of rod saturation relative to M-opsin inputs (i.e. %Rods) when exposed to a commercial display, we made radiometric measurements of a CRT. The CRT’s spectral radiance was compared to rod sensitivity functions (Fig. 8a, red) to calculate R*/rod/s as a function of CRT luminance (Fig. 8b). Then, each value of R*/rod/s was inserted into the model fit in Figure 8d (previously shown with data in Fig. 4c) to yield the fraction of rod input, %Rods. Together, this gave curves of %Rods vs. luminance (Fig. 8e), which is a more relatable domain in most areas of vision science. A typical display produces luminance around 10^2^ cd/m2. At 10^2^ cd/m2, Figure 8b shows that V1 is mostly rod-mediated unless the pupil is fully dilated. This differs from the primate in that 10^2^ cd/m2 is expected to put the retina into a photopic regime. These predictions of %Rods vs. luminance from CRT measurements are expected to generalize across other commercial displays since the shape of their spectral power is similar. Finally, we reiterate that these CRT predictions only pertain to M-opsin in the cone mosaic, which are expressed only in the dorsal half of the retina. The spectral power from commercial displays does not overlap with the spectral sensitivity function of S-opsin, so the cone mosaic will remain silent in the ventral half of the retina, regardless of background intensity.

**Figure 8.**
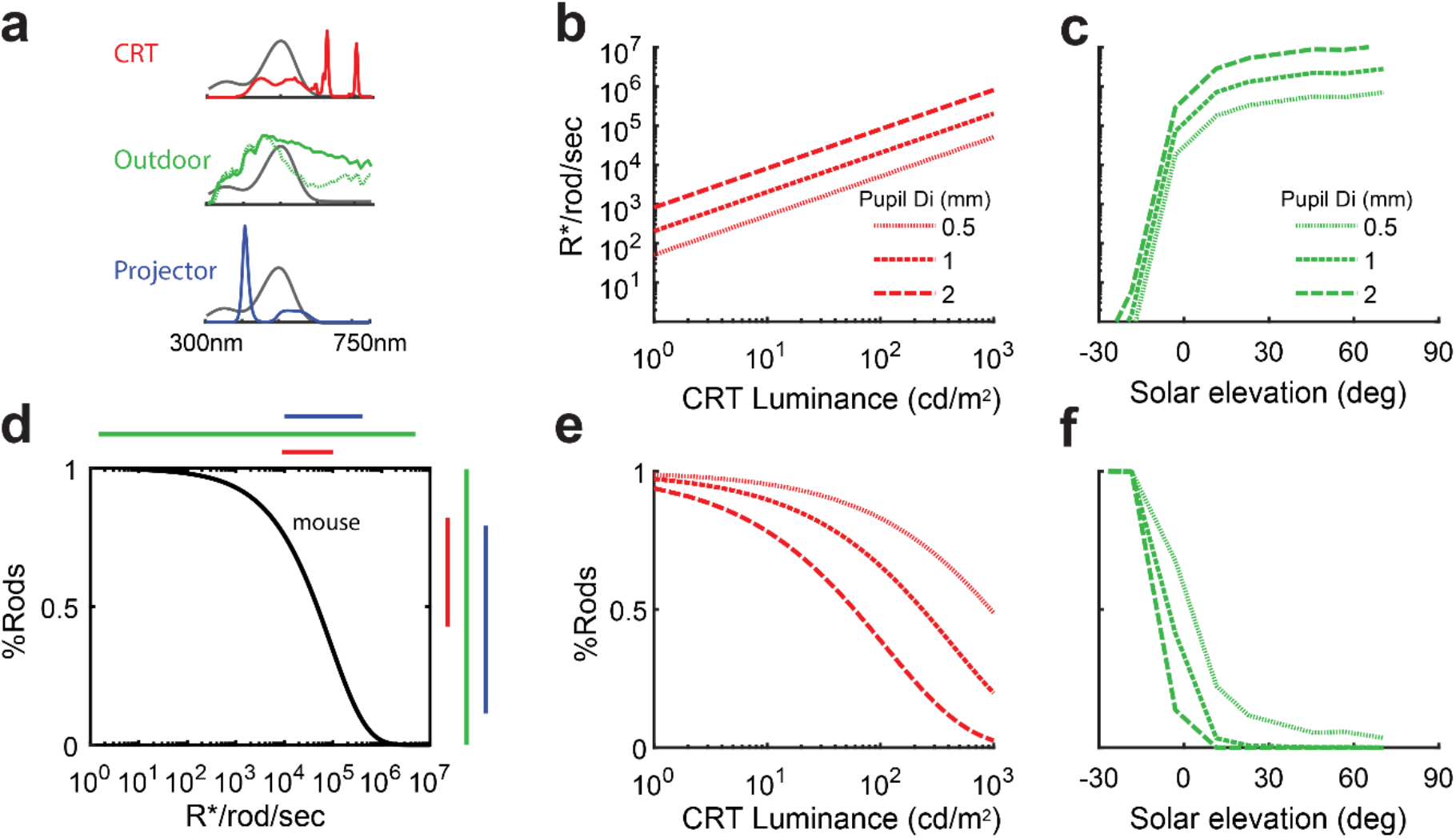
Modeling photoreceptor contributions in the mouse retina based on natural and laboratory stimuli. (a) Spectral power of three different visual stimuli, overlayed on the rod sensitivity function. In red is the power distribution of a CRT with RGB guns all ‘on’. In solid green is the power distribution of the CIE D65 standard, used as the daylight spectrum. The dashed green line is the spectrum used after sunset. Absolute daylight irradiance values were obtained from Spitschan et al., 2016. In blue is the spectrum of the projector used in this study, with both green and nUV LEDs on. (b) Mouse rod isomerization rates over a wide range of CRT luminance. (c) Rod isomerization rates at varying solar elevation in an urban setting. (d) Model fit of %Rod contribution vs. rod isomerization rates, used to transform the plots in ‘b’ and ‘c’ into the plots in ‘e’ and ‘f’. The color bars on top approximate the range of rod isomerization rates generated by each stimulus source. To the right of the y-axis shows the ability of each source to push the mouse retina into a cone-mediated (%Rod = 1) vs. rod-mediated (%Rod = 0) state. (e) Prediction of %Rod contribution as a function of CRT luminance values, for 3 pupil diameters. (f) Prediction of %rod contribution as a function of solar elevation, for 3 pupil diameters. Positive and negative is above and below the horizon, respectively.

If commercial displays cannot yield a cone-mediated mouse V1, a natural follow-up question is whether daylight can produce cone-mediated V1. For this, we used spectral irradiance measurements 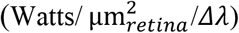 at variable solar elevation taken in a recent study (Fig. 8a, green) (Spitschan et al., 2016). Combined with the rod sensitivity function, the solar spectral irradiance measurements gave R*/rod/sec vs. solar elevation (Fig. 8c). Finally, we plugged R*/rod/sec into our model fit (Fig. 8d) to give %Rod vs. solar elevation (Fig. 8f). Unlike a commercial display, solar radiation has power at wavelengths to drive both M and S opsin. The curves show that the mouse retina rapidly approaches a photopic state after sunrise. Figure 8f provides a summary comparison of the dynamic range of light adaptation in the mouse retina for three lighting conditions. Natural outdoor lighting (green line) spans the full range, from scotopic to photopic. Both of the two experimental displays examined - our projector set-up and commercial displays - place the retina in the mesopic transition zone. However, the relatively incremental advantage provided by the nUV power and overall brightness in our projector system (Fig. 8d; compare blue and red, on top) is able to push the retina near a photopic state (Fig. 8d; compare blue and red, on right). In summary, although we have used seemingly extreme experimental condition to approach photopic vision, the difference from commercial displays is minor within the relatively widespread bounds of outdoor lighting.

A final note on interpretation of these results: the model of %Rod vs. rod isomerization rate is independent of cone adaption under the assumption that cones are above the dark noise, and into Weber adaption. It is for this reason that we only need to consider overlap with the rod sensitivity function to compute %Rod at a given level of light adaptation. For instance, our nUV LED does not have much overlap with S-opsin sensitivity (Fig. 1a,d), yet has sufficient power to place the S-cones into Weber adaption at the lowest light levels studied (Fig. 4a,b). In the case of the CRT, there is no overlap with S-opsin, so these plotted predictions only pertain to M-opsin in lower visual fields. In upper fields, the %Rod prediction is 100% with a commercial display.

## DISCUSSION

The visual stimuli needed to drive cones in and around the fovea is well understood in the primate, which has guided the engineering of commercial displays that drive the perceptual gamut of trichromatic primate vision. The mouse retina, while exhibiting many similarities in structure and function to the primate, lacks a fovea and contains UV-sensitive cone-opsin. Furthermore, the geometry and optical properties of the mouse’s eye produce retinal irradiance for a given stimulus that will be different than in the primate eye. Overall, the balance of inputs from rods and cones to the rest of the mouse’s visual system is difficult to extrapolate from primate psychophysics and rodent recordings in the retina. Here, we used a simple measure of color tuning to model the balance of rod and cone inputs to mouse V1, as a function of background light levels. With this model in-hand, we simulated the balance of rod and cone input to V1 when the eye is exposed to other relevant environments – sunrise puts the mouse retina into a photopic regime, whereas a commercial display does not. Finally, we used our calibration of photoreceptor inputs to design experiments that test the differential contributions of rods and cones to spatio-temporal tuning in V1. Using two different experimental methods, we show a significant increase in responsiveness to higher temporal frequencies in the transition from rod- to cone-mediated vision. We then used the same methods to show that cone-mediated vision produces narrower receptive fields, yet comparable spatial frequency tuning.

### Photoreceptor drive with commercial displays in prior studies of mouse visual cortex

Prior studies of the mouse visual cortex have largely ignored the photoreceptor classes being driven. Nevertheless, one can assume that visible wavelengths in the upper visual field produce rod-mediated responses. In the lower visual fields, the cones express green-sensitive M-opsin, and thus are capable of being driven by commercial displays. However, the simulation in Figure 8e shows that the luminance values required for cone M-opsin mediated vision are just outside the range of commercial displays, even with an artificially dilated pupil.

An additional factor to consider with commercial displays is their attenuation of intensity with increasing viewing angle. This is an issue when imaging mouse visual cortex since a wider range of viewing angles are required to drive all the recorded cells across the retinotopy. The %Rod values shown in Figure 8e, for example, are expected to be even higher at the edges of the screen, assuming calibrations were taken orthogonal to the screen. As shown in Fig. 1d, the rear projection display used in this study is quite Lambertian – the spectral radiance does not change with viewing angle. Similar measurements were made of an LCD panel at variable viewing angle. It was found that peak radiance off the LCD panel was attenuated by 45% and 95% at 30° and 60° viewing angles, relative to the max at 0° viewing angle.

### The required light levels to reach cone-mediated responses can vary

The light intensity required to saturate rods has been measured at varying stages along the rodent visual pathway, and via medley techniques (Naarendorp et al., 2010; Nakatani et al., 1991; Soucy et al., 1998; Wang et al., 2011; Yin et al., 2006), placing it somewhere between 10^3^ and 10^5^ R*/rod/sec. Although several factors are likely to contribute to this variability, a recent and notable study showed that a critical variable is the length of adaptation time (Tikidji-Hamburyan et al., 2017). After 10 min of exposure to 10^5^ R*/rod/sec, an intensity generally presumed to saturate rods, they showed that ~50% of ganglion cells regain their responsiveness. Our brightest light levels resulted in cone-mediated responses following a 10 min adaptation window, which may seem contradictory to these reported temporal dynamics of rod recovery. However, there are two distinctions worth noting.

First, our brightest light levels are just above 10^5^ R*/rod/sec, where rod saturation appears to be more sustainable (Tikidji-Hamburyan et al., 2017; Wang and Kefalov, 2009; Wang et al., 2011). Second, we calculated the balance of rods vs. cones (%Rods), not absolute rod saturation. Residual rod signals may indeed remain at our brightest adaptation point (‘b6’), yet are very small relative to the cones. Furthermore, accumulated noise at the level of cortex precludes the detection smaller signals in the retina. Finally, there is the question of rod recover dynamics at the lower light levels in our adaptation paradigm (b1 - b5). These were played out in descending order following the brightest background condition (b6), and thus expected to be in a steadier regime of rod recovery’s temporal dynamics (Tikidji-Hamburyan et al., 2017).

Differences in rod saturation across studies may be further accounted for by two related experimental variables: 1) presence or absence of rod-cone interactions in the retina, and 2) recording location along the retinotopy. The overlapping spectral sensitivities of rods and cones makes it challenging to drive them independently. For this reason, studies have used mouse lines that lack functional cones. However, mice lacking cone function to assess rod saturation will not be subject to the substantial cone-rod crosstalk in the mouse (Demb and Singer, 2012; Grimes et al., 2018). Furthermore, more recent studies have shown that photoreceptor distributions and retinal circuits are more demarcated along the dorsoventral axis than originally thought (Joesch and Meister, 2016; Nadal-Nicolás et al.; Szatko et al., 2020). As a result, rod-cone interactions may also affect rod saturation in mice with normal retinas, in a retinotopic-dependent fashion.

### Rod- vs. cone-mediated temporal receptive field properties in mouse visual cortex

Cone-mediated temporal dynamics in the retina are much faster than rods, and retinal ganglion cells respond to higher temporal frequencies when inputs originate from cones (Wang et al., 2011). Therefore, a cone-induced tuning shift toward higher temporal frequencies by most V1 neurons (Fig. 5) is expected based on inheritance from responses at the output of the retina. However, the magnitude of the tuning shift in V1 cannot be directly inferred from retinal studies, as the cortex is known to attenuate higher temporal frequencies – there is low-pass filtering at the first synapse from LGN inputs (Freeman et al., 2002), but also between layer 4 and our recording location, layer 2/3 (Niell and Stryker, 2008). This is consistent with the fact that the shift we observed in V1 is not quite as dramatic as what was observed in retinal ganglion cells by (Wang et al., 2011). Another factor that may have influenced temporal frequency tuning in our preparation was anesthesia. V1 response dynamics in awake mice have much more rapid dynamics than anesthetized mice (Haider et al., 2013), which may translate to greater responsiveness to higher temporal frequencies. Therefore, the cortical attenuation of rapid LGN dynamics may be alleviated in the awake prep, thus leading to a more dramatic shift to high temporal frequencies with cone-mediated vision than what has been observed in Figure 5.

### Rod- vs. cone-mediated spatial receptive field properties in a rod dominated retina

The receptive fields of RGC are larger under rod-than cone-mediated light levels, (Cowan et al., 2017; Grimes et al., 2014), and it is expected that V1 neurons would inherit this modulation. Indeed, we found that V1 receptive fields exhibit a 20% increase from cone to rod stimuli (Fig. 7). On the other hand, spatial frequency tuning was practically indistinguishable between rod and cone stimuli. One might expect that spatial frequency and size would exhibit anti-correlated variation, yet our observations are consistent with a simple feedforward model of geniculate inputs to V1 (Hubel and Wiesel, 1962), whereby ON and OFF geniculate afferents converge onto a V1 cell to create a multiphasic ON-OFF receptive field. In this classic model, modest changes in the size of the ON and OFF receptive fields have minimal effect on the ON-OFF separation, and thus peak spatial frequency. Furthermore, similar V1 spatial frequency tuning across rod- and cone-mediated inputs is consistent with human psychophysics in the rod-dominant periphery which show that acuity changes little across the mesopic range of background luminance levels (Hess et al., 1987; Kerr, 1971).

## Methods

### Animal preparation for surgery and imaging

All experiments were approved by the University of Texas at Austin’s Institutional Animal Care and Use Committee. Mice of either sex from C57BL/6 and 129SV strains were used: 43 wild-type mice (C57BL/6; aged 2-5 months; 25 male), and 27 Gnat1^−/−^ mice (129SV; aged 2-5 months; 15 male). The Gnat1^−/−^ strain has a mutation in the rhodopsin gene, causing rod dysfunction. The Gnat1^−/−^ was originally generated on the BALB/c background (Calvert et al., 2000) in the laboratory of Dr. Janice Lem (Tufts University, Medford, MA) and was donated for this study by Dr. Alapakkam P. Sampath (University of California, Los Angeles, CA, USA).

For all surgical procedures, mice were anesthetized with isoflurane (3% induction, 1-1.5% surgery), and given a pre-operative subcutaneous injection of analgesia (5mg/kg Carprofen) and anti-inflammatory agent (Dexamethasone, 1%). Mice were kept warm with a water-regulated pump pad. Each mouse underwent two surgical procedures. The first was the mounting of a metal frame over the visual cortex using dental acrylic, which allowed for mechanical stability during imaging. The second was a craniotomy (4-4.5mm in diameter) over V1 in the right hemisphere, virus injections, and a window implant (3-4mm glass window). Surgical procedures were always performed on a separate day from functional imaging. On a day between the frame implant and virus injections, we identified the outline of the V1 border by measuring the retinotopic map with intrinsic signal imaging (ISI) through the intact bone (Juavinett et al., 2017; Rhim et al., 2017). This allowed for a more uniform distribution of virus across the V1 topography

The virus (pAAV.Syn.GCaMP6f.WPRE.SV40, Addgene viral prep # 100837-AAV1) (Chen et al., 2013b) was delivered using a Picospritzer III (Parker) or Nanoliter injector (WPI) to 2-3 sites in V1 with 0.25-0.5ul per site. Once the injections were complete, craniotomies were sealed using Vetbond and dental cement with dual-layered windows made from coverglass (4mm glued to 5mm, Warner Instruments), and covered between imaging sessions.

Common to all functional imaging procedures, isoflurane levels were titrated to maintain a lightly anesthetized state while the mouse laid on a water-circulating heat pad. However, mapping the V1 retinotopy using ISI is typically more tolerant to high isofluorane levels. For ISI, isoflurane levels were set to 0.25-0.8% (typically 0.5%). For two-photon and widefield calcium imaging, each mouse was given chlorprothixene (1.25 mg/kg) intramuscularly and isoflurane levels were adjusted to 0.25-0.5%. Silicone oil (12,500 cst) was periodically applied to the stimulated (left) eye to prevent any dryness or damage to the eye. Silicone oil allows optical transmission of near-UV (nUV) and visible wavelengths. Additionally, the unstimulated (right) eye was coated with Vaseline and covered with black tape during imaging.

### Imaging

Widefield images were captured using a Pco Panda 4.2 camera with Matlab’s image acquisition toolbox, at 10-to-20 frames/sec, and 2×2 binning. An X-Cite 110LED (Excelitas) was used for illumination. For intrinsic signal imaging, a longpass colored glass filter (590nm) was placed over the light guides to illuminate the brain, and a bandpass filter (650 +/− 25 nm) was on the camera lens. For widefield fluorescence GCaMP6f imaging, a GFP filter cube was placed on the camera lens.

Two-photon calcium imaging was performed with a Neurolabware microscope and Scanbox acquisition software. The scan rate varied between 10-15 frames/sec, scaling with the number of rows in the field-of-view. A Chameleon Ultra laser was set to 920 nm to excite GCaMP6f. A Nikon 16x (0.8NA 3mm WD) or Olympus 10x (0.6NA 3mm WD) objective lens was used for all imaging sessions.

### Visual stimuli

#### Setup overview

Two monochrome LED projectors (Keynote Photonics) with spectral peaks at 405 nm (nUV) and 525 nm (green) were used to generate spatio-temporally modulated stimuli that drive opsins in the mouse retina (Fig. 1a-d) (Rhim et al., 2017). The refresh rate was 60 Hz. The rear projector screen was made of Teflon, which provided a near-Lambertian surface (Fig. 1d). The two projectors were first independently mounted on an articulating platform to align the images as closely as possible, followed by a more refined software alignment consisting of a lateral shift and affine transformation. This software calibration procedure first entailed showing a grid of dots by the green projector, and a user-controlled mouse shown by the nUV projector. The user is prompted to “click” on each point in the grid, which is used to compute the transformation that aligns the projector images. Stimuli were coded using the Psychophysics Toolbox extension for Matlab (Brainard, 1997; Pelli, 1997)

The mouse was positioned so that its left eye was vertically centered on the projector screen, and the perpendicular bisector from the mouse’s eye to the screen was between 8 and 10 cm. Next, the screen was angled at 30° from the mouse’s midline. The final constraint was to align the front edge of the screen to the mouse’s midline.

Image size and intensity vary with the distance between the projector and the Teflon screen. We selected a screen-to-projector distance that yielded a maximum image size just large enough to drive the majority of V1, while keeping intensity high. The screen size was 43 cm high × 23.5 cm wide, which gives approximately 135° × 105° of visual angle.

#### Measuring the uniformity of spectral radiance

Our experiments required visual stimuli that are calibrated for UV and visible wavelengths across a wide range of viewing angles, which ultimately requires an efficient diffuser of rear-projected nUV and visible light. We tested multiple commercial rear-projection films, but found that 0.01” Teflon (McMaster-Carr) (Tan et al., 2015) was unmatched in its optimization of our criteria. Criteria were based on uniformity of spectral radiance, across screen position and viewing angle. For each measurement, both green and nUV projectors showed a uniform image on the opposite side of the rear projection film as a PR655 spectroradiometer. The spectroradiometer was focused on the screen at 7 locations - center, top edge, bottom edge, and each corner (only center location measurement shown in Figure 1d). At each location, a measurement was taken at three viewing angles relative to the planar screen, 30°, 60°, and 90°, totaling 21 measurements of spectral radiance. Ideally, the spectral radiance does not change across these 21 conditions. Figure 1d summarizes the results from the material we settled on (0.01” Teflon) for the center location. The spectral radiance curves are virtually identical across viewing angles. Furthermore, the amplitude of the radiance curve across the various other locations and viewing angles (not shown) is also quite consistent. This uniformity in spectral radiance is far superior to our measurements of other commercial displays (a CRT and LCD), other commercial rear projection materials, and thinner Teflon (0.005”). We would expect that the uniformity to be maintained for thicker Teflon, yet this would have the undesirable effect of attenuating the overall intensity of the image.

#### Stimulus A: Retinotopic mapping stimulus

The same drifting bar stimulus was used to map retinotopy, regardless of imaging modality. This method is an extension of the one used in (Kalatsky and Stryker, 2003), and described in detail in (Marshel et al., 2011). To summarize, a periodic drifting bar on black background was modulated by a contrast-reversing checkerboard, and shown in four cardinal directions. The bar’s speed and width were dependent on screen location such that the temporal phase of cortical responses, at the stimulus frequency, could be directly mapped to altitude and azimuth coordinates in a spherical coordinate system. The speed of the bar was ~5°/s and its width was ~12°. The checkerboard squares were 5° wide and reversed contrast at a rate of 2 Hz. Both green and nUV projectors were shown at maximum contrast to produce the checkerboard. Each drift direction was shown at ~0.055 HZ for wide-field imaging and ~0.042 HZ for the 2-photon calcium imaging setup. There were 2-to-3 120 sec trials, for each drift direction.

#### Stimulus B: Color tuning at graded light adaptation

With the goal of systematically varying rod saturation, a total of 6 different mean light levels were used across the experiments in this study (b1-b6), while keeping contrast the same (Figs. 1e,2a). Each intensity was fixed for an entire block of drifting grating trials; prior to each, the eye was adapted for 10 min with a uniform screen of matched mid-point intensity. To quantify the graded transition from rod- to cone-mediated inputs, all 6 light levels were used, and the responses to M-isolating and S-isolating gratings (see below) were compared (Figs. 2–4). Furthermore, the spatial and temporal frequencies were kept constant across trials at 0.05 cyc/° and 1 cyc/sec, respectively, which are well represented in mouse V1 (Gao et al., 2010; Niell and Stryker, 2008). Drift direction varied across trials: 0, 45, 90, 135, 180, 225, 270, and 315°. Each stimulus was shown for 3s, flanked by 1s of pre-stimulus and 1s of post-stimulus blanks. The mid-point “gray level” was held constant throughout the entire adaptation and test stimuli.

#### S- and M-opsin isolating gratings

Green and nUV LED sinewaves were combined to produce contrast along the S- and M-isolating directions of cone-opsin space (Fig. 1f). As shown below, S-opsin and M-opsin contrast can be described as a function of the rear-projected spectral radiance of the nUV (*R*_*UV*_(λ)) and Green (*R*_*G*_(λ)) LEDs, the amplitude of their drifting sinewaves (*a*_*UV*_, *a*_*G*_), and the S- and M-opsin sensitivity functions (*h*_*S*_(λ) and *h*_*M*_(λ)):

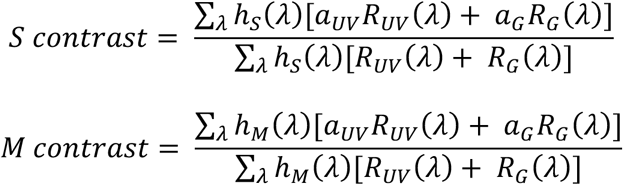

Spectral radiance was measured using a PhotoResearch PR655. Opsin sensitivity functions are taken from (Govardovskii et al., 2000). To null either S- or M-opsin contrast, we utilized “silent substitution” (Estévez and Spekreijse, 1982). M-isolating stimuli (“*S contrast*” = 0) can be achieved by limiting modulation to the Green LED (*a*_*G*_ > 0), while keeping the nUV LED constant (*a*_*UV*_ = 0). This is because *R*_*G*_(λ) and *h*_*S*_(λ) do not overlap (Fig. 1a,d). However, for S-isolating stimuli (“*M contrast*” = 0), we used the equations above to solve for *a*_*UV*_ and *a*_*G*_. The maximum possible S-opsin contrast for the S-isolating stimulus was 60%. Therefore, the M-isolating stimulus was also shown at 60% contrast. The same calibration procedure was used to generate M and S contrast for *Stimulus D* and *E,* described below.

#### Stimulus C: Spatio-temporal frequency tuning, at low and high light adaptation

Full-field drifting sinewave gratings at variable spatio-temporal frequency settings were used to study the spatio-temporal frequency characteristics of rod-dominated versus cone-dominated regimes in V1. To measure temporal frequency tuning, five temporal frequencies were shown: 0.5, 1, 2, 4, and 8 (cyc/s), all at a spatial frequency of 0.05 cyc/deg (Fig. 5a-f). Only one color was shown, which had 60% contrast along the S+M axis of cone-opsin space (Fig. 1f). One of eight drift directions, 45° apart, were shown on each trial. Each stimulus in the ensemble was shown 5 times, giving the following number of trials: 5 × 8 × 5 (*temporal frequency* × *drift direction* × *repeat*; 200 trials). Each stimulus was shown for 3s, flanked by 1s of pre-stimulus and 1s post-stimulus blanks. This 200-trial stimulus block was shown at two different states of light adaptation, “b1” (dimmest) and “b5” (2^nd^ brightest). Each block was preceded by 10 minutes of adaptation at the same gray level. Next, the spatial frequency stimulus used the same template – two 200-trial light adaptation blocks that varied spatial frequency as follows: 0.0125, 0.025, 0.05, 0.1, 0.2 cyc/deg (Fig. 6a-h). Temporal frequency was held at 1 cyc/sec.

#### Stimulus D: Spatio-temporal frequency tuning, with rod- and cone-isolating contrast

Here, spatial or temporal frequency tuning was measured in a single mesopic adaptation block that varied opsin contrast. In the temporal frequency version of this experiment (Fig. 5g-i), stimuli had one of 5 temporal frequencies [0.5 1 2 4 8 cyc/sec], 2 color contrasts [M- or S-isolating], 8 drift directions (45° intervals), and 1 spatial frequency (0.05 cyc/deg). In the spatial frequency version (Fig. 6i-l), stimuli had one of 5 spatial frequencies [0.0125 0.025 0.05 0.1 0.2 cyc/deg], 2 color contrasts [M- or S-isolating], 8 drift directions (45° intervals), and 1 temporal frequency (1 cyc/s). In each case, light levels were held constant at either ‘b5’ or ‘b4’, which place the retina in a mesopic state. Also, the stimulus referred to as “M-contrast” (above) is a rod-isolating contrast for neurons in the upper visual field in a mesopic state.

#### Stimulus E: Mapping receptive fields, with rod- and cone-isolating contrast

To measure the width of receptive fields along the vertical axis of visual space (Fig. 7), horizontal bars were flashed on the screen at varying vertical position and color. The four possible color directions of the bar in the S/M opsin plane were S-, S+ M-, and M+. In the upper visual field, M-opsin contrast only drives rhodopsin. The bars were 6° wide in visual angle, had a square spatial profile, and were sampled every 2.8°. Each bar was shown, static, for 0.75s with pre- and post-blank period of 0.25s each. The horizontal length of each bar extended from the left to right edge of the screen. Each stimulus combination of color and position was repeated 20-30 times, and signals were averaged for each.

### Eye dilation experiment and rod isomerization rate

In a subset of experiments, *Stimulus B* (above) was run before and after pharmacologically-induced pupil dilation (1% tropicamide) on the same day. To track the pupil diameter across experiments a CMOS camera with IR sensitivity (Allied Vision Manta G-235B) was mounted with a long working distance Zoom lens (Navitar 6000) and a 590 nm longpass filter. The camera and lens were just behind the mouse and pointed at a 5mm angled mirror within the mouse’s field-of-view to capture the pupil images. During 2-photon imaging, the IR excitation transmitted through the head and out of the eye for a crisp image of pupil diameter (Fig. 2b). After each block of stimuli set to a given level of light intensity, a picture of the pupil was taken. Pupil diameter was measured along the axis that roughly appeared to yield the maximum length since the image was taken at an angle.

### Quantifying photoreceptor isomerization rates for each light level

To estimate the photoisomerization rates of rods and cones in our preparation, we start by converting the measured radiance from the display, in 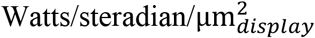, into irradiance at the retina, in 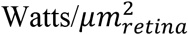. The first step is to multiply the display radiance by the steradians at the pupil. Steradians is roughly the area of the pupil over the squared distance between the display and the retina, A_pupil_/D^2^_*display*_. Next, to convert area on the display,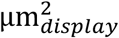, to area on the retina, 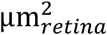, we divide by the squared magnification of the image formation, D^*2*^_*retina*_/D^*2*^_*display*_, where D_*retina*_ is the diameter of the retina and D_*display*_ is the distance to the display. To simplify, D^2^_*display*_ cancels out, and we can just multiply the original radiance measurement by A_pupil_/D^*2*^_*retina*_ to get irradiance, in 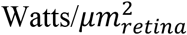. The irradiance values are sampled at discrete wavelengths, Δλ, which gives units of 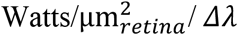. Finally, we scale by the transmittance at each wavelength, *T*(λ), to compute the final retinal irradiance. We used a model for *T*(λ) through the mouse lens, given in (Lei and Yao, 2006), which attenuates by 22.5% at 380 nm, and drops monotonically to 8.0% attenuation at 700 nm. To formalize, we convert radiance off the display, ′*Rad*(λ)′, into irradiance at the retina, ′*I*(λ)′ as follows:

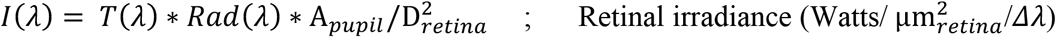

Next, convert Joules to quanta

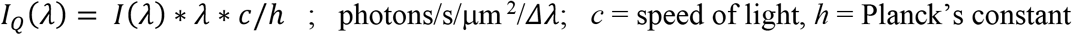

To compute isomerization rate, *R*∗/*receptor*/*S*, we take the dot-product between the quantal retinal irradiance, *I*_*Q*_(λ), and the absorption spectrum, *a*_*c*_(λ), and scale by the sample period of the spectrum Δλ:

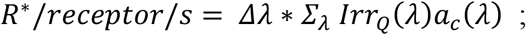

To get *a*_*c*_(λ) in μm^2^, we started with the unitless absorption spectra of rods, M-opsin, and S-opsin from (Govardovskii et al., 2000) and then scaled by the end-on collection area at the peak wavelength which may be approximated as near 1 and 0.85 μm^2^ for cones and rods, respectively (Naarendorp et al., 2010). To get the final isomerization rate in a given experiment, we scale by the corresponding projector intensity (e.g. buffer value), relative to the values obtained from the initial measurement of *Rad*(λ). This gives the following range of isomerization rates, across the 6 light adaptation levels (Figure 2a), for each opsin: M-opsin = [43.0K 430K], S-opsin = [5.7K 57K], and rhodopsin = [12K 382K].

### Quantifying photoreceptor isomerization rates as a function of solar elevation

A recent study measured solar irradiance in a rural setting, in absolute power units, with varying solar elevation (Spitschan et al., 2016). Their calibrated units in Watts/m^2^ allow us to quantify isomerization rates in the mouse retina. For the nighttime irradiance, we used the same spectral profiles given in Spitschan et. al., as these contained values into UV. However, their daylight measurements did not contain the UV part of the spectrum, which is important for computing isomerization rates in the rodent. In turn, we used the unitless CIE D65 standard for daylight irradiance, which does include UV values, and scaled it by the Spitschan et. al. irradiance measurements at 450nm. Spitschan et. al. found that the biggest deviations from the CIE D65 standard occur at lower solar elevations, which validates the use of the CIE D65 standard at higher solar elevations. In the equation below, *I*_*outdoor*_(λ|∅) is the spectral irradiance in W/m^2^ at discrete solar elevations, ∅. To obtain the spectrum at each ∅, we took the average within a range of elevations, and across the medley variables that can affect it on a given day (Spitschan et al., 2016). Next, the outdoor irradiance was converted into radiance at the retina by multiplying by the surface area of the pupil, A_pupil_, over the surface area of the retina, A_retina_ (Lyubarsky et al., 2004).

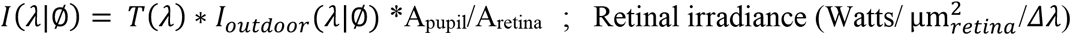

Then, retinal irradiance is converted into isomerization rates, using the same manipulations as above for the display calculations. But now, it is expressed as a function of solar elevation:

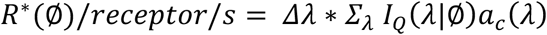

In Figure 8c, this is plotted for the case of *a*_*c*_(λ) being the rhodopsin sensitivity function, at three values of A_pupil_.

### Data analysis

#### Widefield imaging

Widefield images were captured to measure responses to two stimuli. The first was retinotopy (*Stimulus A*). The preferred phase at each pixel, at the frequency of the drifting bar, was computed for each of the four drift directions. The opposing directions (e.g. up and down) were then combined to yield retinotopic position as described in (Kalatsky and Stryker, 2003). This gave a clear boundary of V1 that could be manually outlined. The second stimulus with widefield was used to measure color tuning at graded light adaptation (*Stimulus B*). On each trial, the time course of each pixel was normalized by a 400 ms blank period preceding grating onset, to yield a time course of ΔF/F (i.e. % response above the preceding baseline). Next, the mean was taken from 100 ms to 3100 ms after stimulus onset of the 3000 ms stimulus. We then averaged over 8 drift directions and 5 repeats to get the mean response of each pixel to the M-isolating stimulus and S-isolating stimulus. Finally, the map of *M*_*pref*_ was computed as described in Eq. 1 below.

#### Generating neuronal tuning curves from two-photon movies

Neurons were identified using the local cross-correlation image, whereby the time course of each pixel was cross correlated with the weighted sum of its neighbors. This is given as

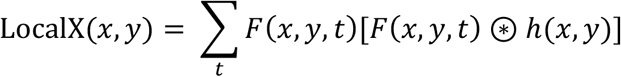

where *h* is a Gaussian with σ = 3 microns, and ⊛ indicates convolution in *x* and *y*. ‘LocalX’ contains high contrast puncta indicating regions of high temporal correlation that are more independent from the background. A very similar procedure was performed in a prior study (Smith and Häusser, 2010). The puncta were then manually selected using a custom “point-and-click” GUI created in Matlab. The pixels were then averaged to give the time course of each cell.

On each trial, a neuron’s raw fluorescence timecourse was converted into units of *ΔF/F*. Specifically, *ΔF/F* = [*F*(*t*)-*p*]/*p*, where *F*(*t*) is the raw signal and *p* is the mean fluorescence during a blank screen preceding the trial. The mean of *ΔF/F* was then taken during the response. Most stimuli used in this study consisted of trials with drifting sine wave gratings presented for 3 sec (*Stimulus B-D*), in which case *p* was taken over a 1000 ms window, and the response mean was taken between 500 ms to 3250 ms after stimulus onset. In the case of the static flashed bars (*Stimulus E*), the window for *p* was 250 ms, and the trial integral was taken between 250 ms and 850 ms. For *Stimulus A*, *ΔF/F* was not calculated since we were only interested in the phase.

#### Model of photoreceptor inputs vs. light intensity and retinotopy

At each of 6 background light levels (see *Stimulus B*), we computed each neuron’s color preference metric defined below, where *F*_*S*_ and *F*_*m*_ are the mean *ΔF/F* response to the S- and M-isolating stimuli.

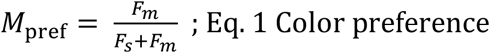

In general, the rods are not fully saturated, so a neuron’s response to each color direction, *F*_*S*_ and *F*_*m*_, is a function of input from both rods and cones, which we represent as a linear combination below.

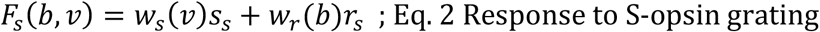

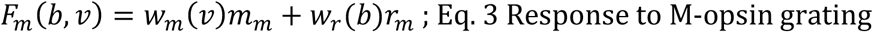

*w*_*S*_(*v*) and *w*_*m*_(*v*) are cone S- an M-opsin input weight as a function of vertical retinotopy, *v*. *w*_*r*_(*b*) is the rod input weight, as a function of rod isomerization rate, 10^3^ *R*∗/*rod*/*S*, denoted *b*. Each opsin weighting function is unitless, but scaled relative to each other. For instance, under low light levels where rods dominate, then *w*_*r*_>>*w*_*s*_. Also, the sum of cone opsin inputs is unity across the retinotopy; i.e. *w*_*s*_(*v*) = 1 − *w*_*m*_(*v*). To get the response contribution of each opsin, the weighting function is scaled by a stimulus contrast; *S*_*s*_ is the S-opsin contrast for the S-isolating stimulus (~0.6), and *m*_*m*_ is the M-opsin contrast of the M-isolating stimulus (~0.6). *r*_*s*_ and *r*_*m*_ are the rod contrasts to the S- and M-isolating stimuli. Plugging Eqs. 2 and 3 into Eq. 1 gives

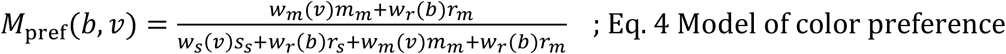

Unlike rod weights in this model, cone weights do not depend on the light level. This is based on our measurements in Gnat1^−/−^ mice, where responses were roughly constant across the range of light levels, *b* (Fig. 4a-b). Also, the lowest light levels in our experiments place the isomerization rate of each cone just above 5K R*/cone/s, which is above the dark noise according to prior studies, at which point Weber adaptation takes over (Schnapf et al., 1990; Schneeweis and Schnapf, 1999; Naarendorp et al., 2010). Lastly, whereas rod weights are assumed independent of the retinotopy since the mouse lacks a fovea, the balance of M and S-opsin weights are dependent on the vertical retinotopy (Rhim et al., 2017).

Finally, Equation 4 can be simplified to help with subsequent manipulations and model fitting. Since rhodopsin and M-opsin have very similar sensitivity functions, we can assume that rod contrast to the M-isolating stimulus is much greater than rod contrast to the S-isolating stimulus. Or *r*_M_ ≫ *r*_J_. After plugging in *S*_*s*_ = *m*_*m*_ (i.e. equated cone-opsin contrast) and *w*_M_(*v*) = 1 − *w*_J_(*v*), this simplifies to

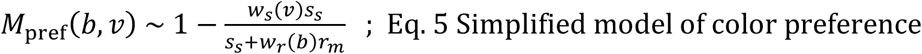

Near rod saturating light levels, *w*_*r*_(*b*) << *w*_*s*_(*v*), so that *M*_*pref*_(*b*, *v*) approaches 1 − *w*_*s*_(*v*) = *w*_*m*_(*v*). At the opposite extreme, when light levels are low, *w*_*b*_(*b*) >> *w*_*s*_(*v*), and *M*_*pref*_(*b*, *v*) approaches 1. *M*_*pref*_(*b*, *v*) is the measured variable and *w*_J_(*v*) and *w*_H_(*b*) are unknowns. Eq. 5b has the convenient feature of separability along *v* and *b*. As described below, we first fit the data along *b*, and then *v*.

#### Calculating rod input vs. R*/rod/sec

To simplify the calculation of rod input, *w*_*r*_(*b*), we limited data points to the upper visual field >30°) where the S-opsin contribution from the cones is near 100% (Wang et al., 2011); i.e. *w*_*s*_(*v* > 30°) = 1.

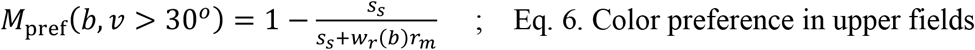

We then fit 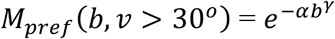 to the data, which gave α = 0.072 and γ = 0.586. Again, ‘*b*’ is in units of 10^3^ R*/rod/sec. Setting 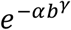 equal to Eq. 6, then plugging in the rod and cone opsin contrast values, *r*_*m*_ ~ *S*_*s*_ = 0.6, and finally solving for *w*_*r*_(*b*) gives

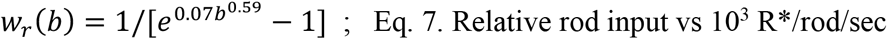

*w*_*r*_(*b*) is unitless and scaled relative to the cone opsin weights. A more meaningful model parameter is obtained by calculating the rod weight, relative to the total rod and cone weight:

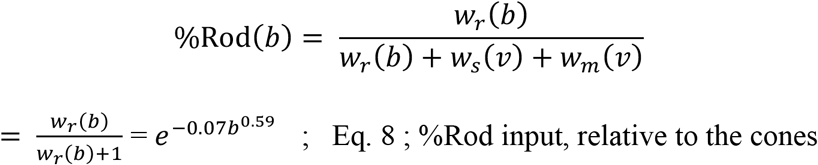

Importantly, Eq. 6 and Eq. 8 are equivalent: *M*_*pref*_(*b*, *v* > 30°) = %Rod(*b*). That is, the measured color preference in the upper visual field is also a measure of the rod preference. Also, we have expressed %Rod(*b*, *v* > 30) as %Rod(*b*), which assumes that rod saturation is constant if irradiance is constant.

#### Calculating cone opsin balance vs. retinotopy

Finally, we fit a model to *w*_*m*_(*v*) = 1 − *w*_*s*_(*v*), which is the balance of M- and S-cone opsin at each location of the vertical retinotopy, independent of the rods. Coming back to Equation 5b, which is the model of the observed variable *M*_*pref*_(*b*, *v*), we can solve for *w*_*m*_(*v*) at each data point since *w*_*r*_(*b*) was solved for in Eq. 7. Then, we fit the following sigmoid (Fig. 4f, red).

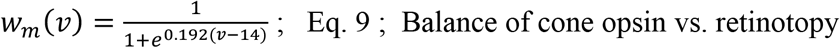

#### Quantifying spatio-temporal tuning

To get the spatial or temporal frequency tuning curve of each cell, at each background level (Stimulus C) or color (Stimulus D), we averaged responses over repeats, and then drift direction. Next, a Gaussian function with a domain of log_2_(spatial frequency) or log_2_(temporal frequency) was fit in order to quantify tuning curves. For both temporal and spatial frequency tuning, center-of-mass and high-pass cutoff were calculated from the fit. For only spatial frequency, the peak location of the fit was also used.

Center-of-mass was determined in the logarithmic domain as CoM_log_ = ∑_*f*_ *G*(log_2_(*f*)) ∗ log_2_(*f*) / ∑_*f*_ *G*(log_2_(*f*)), where *G*() is the Gaussian fit, and *f* is the stimulus domain in cyc/deg or deg/sec. CoM_log_ is not necessarily the peak location of the Gaussian because the sum over *f* is limited to the stimulus domain. Then, CoM_log_ was converted into the same units as *f* by taking 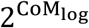. Next, peak location and high-pass cutoff were both determined from a finely sampled version of the Gaussian fit. Note that the peak location differed from the parameter μ in the Gaussian fit when the tuning was low-pass. High-pass cut-off frequency was taken at 70% of the peak’s height, to the right of the peak.

#### Two-photon data yield

All data analysis followed an initial preprocessing pipeline for quality control. To be included in the analysis, each neuron had to cross a response threshold for two stimuli: (1) Its mean ΔF/F response in the stimulus window of the sinewave grating stimulus-of-interest (i.e. *Stimuli B, C, D,* or *E*) and (2) its amplitude of response at the stimulus frequency of the drifting bar stimulus, which was used to map retinotopy (*Stimulus A*). A cell was excluded if (1) it did not have a mean ΔF/F > 0.05 for any stimulus within the ensemble of episodic stimuli, OR (2) if the magnitude of the Fourier amplitude at the stimulus frequency did not have a peak sufficiently greater than the noise – specifically, unreliable activity was identified based on the ratio of the first harmonic (F1) magnitude over the mean of the Fourier magnitudes at neighboring frequencies (between 0 and 0.14 Hz). If this ratio was less than 5 for any drift direction, it was excluded from subsequent analyses.

For experiments that required fitting a Gaussian curve (see“*Quantifying spatio-temporal tuning*”), an additional screening was applied based on the variance accounted by the fit. A threshold of 65% and 45% was applied to WT and Gnat1^−/−^, respectively.

The majority of the animals used in this study generated data for a single figure. Of the 43 wild-type mice, 7 were used for the pupil experiment (Fig. 2), 11 for the graded adaptation experiment (Figs. 3,4), 6 for the temporal frequency experiment (Fig. 5), 14 for the spatial frequency experiment (Fig. 6), and 12 for the receptive field size experiment (Fig. 7). Of the 27 Gnat1^−/−^ mice, 3 were used for the graded light adaptation (Fig. 4a,b), 12 for the temporal frequency experiment (Fig. 5), and 12 for spatial frequency experiment (Fig. 6).

The following summarizes WT cell survivability based on the above threshold criteria applied to the analyses within each figure sub-panel. Pupil experiment (Fig. 2): 386/619 cells. Temporal frequency experiment (Fig. 5a-c): 235/662 cells. (Fig. 5g-i): 218/334. Spatial frequency experiment (Fig. 6a-d): 250/1154 cells. (Fig. 6i-l): 361/818 at 44.13%. Receptive field size mapping (Fig. 7): 108/1082 cells. For the graded light adaptation experiments in 11 WTs (Figs. 3,4), the yield varied slightly across the different states of adaptation, “b1-b6”. In 2 of the 11 animals, b6 was not run. For b6, the yield was 666/1014 cells. For b1-b5 (11 animals), the yield ranged from 773/1252 at b2, up to 791/1252 at b5. The Gnat1^−/−^ yield was as follows: Temporal frequency experiment (Fig. 5): 244/1218 cells at 20.03%. Spatial frequency experiment (Fig. 6): 72/838 cells at 8.59%. For the graded light adaptation (Fig. 4a,b), the yield ranged from 94/215 at b1, up to 102/215 at b6.

## CODE AVAILABILITY

Matlab code contributing to this study is available upon reasonable request.

## DATA AVAILABILITY

Data from this study is available upon reasonable request.

## ACKNOWLEDGEMENTS

This work was supported by the Whitehall Foundation and the NIH R01EY028657

## AUTHOR CONTRIBUTIONS

IN and IR designed the experiments. IR and GCR performed surgical procedures and imaging. IN and IR analyzed the data and wrote the paper. All authors assisted with revisions.

